# The human brain reactivates context-specific past information at event boundaries of naturalistic experiences

**DOI:** 10.1101/2022.06.13.495935

**Authors:** Avital Hahamy, Haim Dubossarsky, Timothy E. J. Behrens

## Abstract

Although we perceive the world in a continuous manner, our experience is partitioned into discrete events. However, to make sense of these events, they must be stitched together into an overarching narrative – a model of unfolding events. It has been proposed that such a stitching process happens in offline neural reactivations when rodents build models of spatial environments. Here we show that, whilst understanding a natural narrative, humans reactivate neural representations of past events. Similar to offline replay, these reactivations occur in hippocampus and default mode network, where reactivations are selective to relevant past events. However, these reactivations occur, not during prolonged offline periods, but at the boundaries between ongoing narrative events. These results, replicated across two datasets, suggest reactivations as a candidate mechanism for binding temporally distant information into a coherent understanding of ongoing experience.

## Introduction

Reading a scientific paper is a complicated task. The paper you are about to read is composed of a stream of words that convey a detailed, interconnected narrative, that you will try to make sense of. Although you may read through this narrative continuously, your brain will process the narrative by parcelling it into discrete chunks of information, termed “events”^1^, with associated neural representations (“event models”^2, 3^). The transitions between events, termed event boundaries, have been suggested to occur when shifts in the content of incoming information will be detected, at which point the current event model will be encoded into memory, and a new event model will be formed^1, 2, 4, 5^. This could happen, for example, in the transition between paragraphs.

However, understanding a narrative requires more than just parcelling the ongoing stream of information into events and storing them into memory. Following an ongoing narrative requires relations to be drawn between each current event and contextually-relevant past events (for example, linking results presented in the Results section with their related research hypothesis presented in the Introduction section). How, then, could these events be pieced together? It has long been proposed that remote past information could be integrated with incoming sensory information^6-8^. Accordingly, it has been suggested that event-models of past events can assist in the interpretation of incoming information^5, 9-12^. However, the mechanism underlying this process is currently unknown.

One possible mechanism that may underlie this integration is termed “replay”, and was originally seen in rodents. Here, sequences of cell firing patterns that are observed during past events are rapidly recapitulated during pauses in behaviour. Replay has predominantly been studied offline, during periods of rest or sleep^13, 14^, but it can be triggered by salient behavioral events, for example when an animal completes a trial^15^ or receives a reward^16^. These situations could be analogous to event-boundaries, as they mark contextual shifts in the ongoing experience. This hypothesis is supported by human imaging studies, demonstrating that the representations of just-ended events are reactivated at event boundaries^17, 18^ (due to the lack of temporal resolution to infer a sequentially-ordered reinstatement, the term reactivation, rather than replay, is commonly used).

Event segmentation and replay also engage similar brain regions. Increased activity at event boundaries during the processing of a narrative has been associated with regions including the hippocampus^19, 20^ and nodes of the default mode network (DMN^21^, mainly the precuneus, angular gyrus/superior lateral occipital cortex and superior/middle temporal gyrus^5, 22-24^). Similarly, replay and sharp wave ripples are associated with increased activity in hippocampus and in the DMN in both humans and animal models^25-29^.

It has been proposed that both replay in rodents and reactivation in humans encode past events into memory, as well as inform future decisions^13, 30^. But, notably, offline replay in rodents is not limited to playing a veridical recording of experience. Instead, it can make inferences - piecing together multiple past events, even remote ones, as if building, and sampling from, a model of the world^13, 31- 33^. Similarly, in humans, replay can reorganize and reorder events to fit with a pre-learned task model^34, 35^. It is therefore possible that, as an experience is unfolding, reactivation could underlie the inference of relations between current and past events, and continuously bind related event-models as the narrative is progressively built.

Replay studies have, so far, used controlled experiments with regularly-repeated trials, which make it hard to assess the integration of events across time, since repeated events would have very similar representations. Yet, the integration of events across time can be measured in more naturalistic experiments that have a continuously evolving narrative: here, each event is unique and should have a more distinct representation. In such experiments, we would expect to find that reactivation at event boundaries would not only contain representations of the event that has just ended^17^. Reactivation should also contain representations of unique past events, particularly those most relevant to the understanding of the current narrative stage (for example, you will gain a better understanding of the current hypothesis, if at the relevant event boundary, your brain replays the hypothesis along with supporting information from previous paragraphs).

We therefore aimed to detect brain regions in which, not only the event that has just ended is reactivated at event boundaries, but also more temporally-remote events that are relevant to the understanding of the present situation. To this end, we analysed two naturalistic datasets, where participants either viewed a movie or listened to a story while undergoing a whole-brain fMRI scan. We examined these datasets using a novel analytic method, designed to detect reactivated representations of remote events at event boundaries. Using this method, we indeed revealed reactivation in the posterior hippocampus, the precuneus/retrosplenial cortex, and the angular gyrus/lateral occipital cortex, and replicated these results across datasets. We further demonstrated that this reactivation is apparent at both a local, voxel-scale, and also at a meso-scale. Finally, we found that the reactivated representations in the precuneus/retrosplenial cortex were of semantically-relevant past events, demonstrating a specific integration of information that is needed for the understanding of each current narrative stage. We therefore propose that reactivation is a constructive mechanism that builds our understanding of evolving experiences by selectively binding relevant pieces of information across time.

## Results

Can reactivation at event boundaries integrate current, ongoing experiences with temporally-remote past information? To answer this question, we used fMRI data acquired from participants engaged in a naturalistic narrative (movie/story). Reactivation entails the expression of past scene-representations (voxel-activity patterns) at event boundaries. Therefore, to detect reactivation in these fMRI data, we looked for correlations between fMRI representations at event boundaries (scene endings) and representations of remote scenes (preceding the just-ended scene) using Representation Similarity Analysis^36^. However, due to the low temporal resolution of fMRI data, representations at event boundaries are likely correlated with the representations of the immediately preceding scenes. If these scenes are similar to other scenes, then correlations between event boundaries and remote scenes may in fact reflect between-scene correlations, rather than reactivation of remote information. This problem can be solved, since between-scenes correlations are symmetric in time (past/future, see Figure 1b). We therefore defined a “reactivation index” that cancels out symmetric similarities by contrasting past with future correlations. The results obtained from this reactivation analysis were replicated across two independent fMRI datasets (Sherlock dataset^37^, 21st year dataset^38^, see Table 1). We demonstrate these results using reactivation within the brains of single participants, and also using the similarity in reactivation between the brains of different participants (potentially measuring reactivation of representations at a mesoscopic scale). Finally, we used the overlap in words between scenes to test whether the reactivated scenes are those that are most relevant (semantically-similar) for the understanding of the current scene.

**Table 1.**
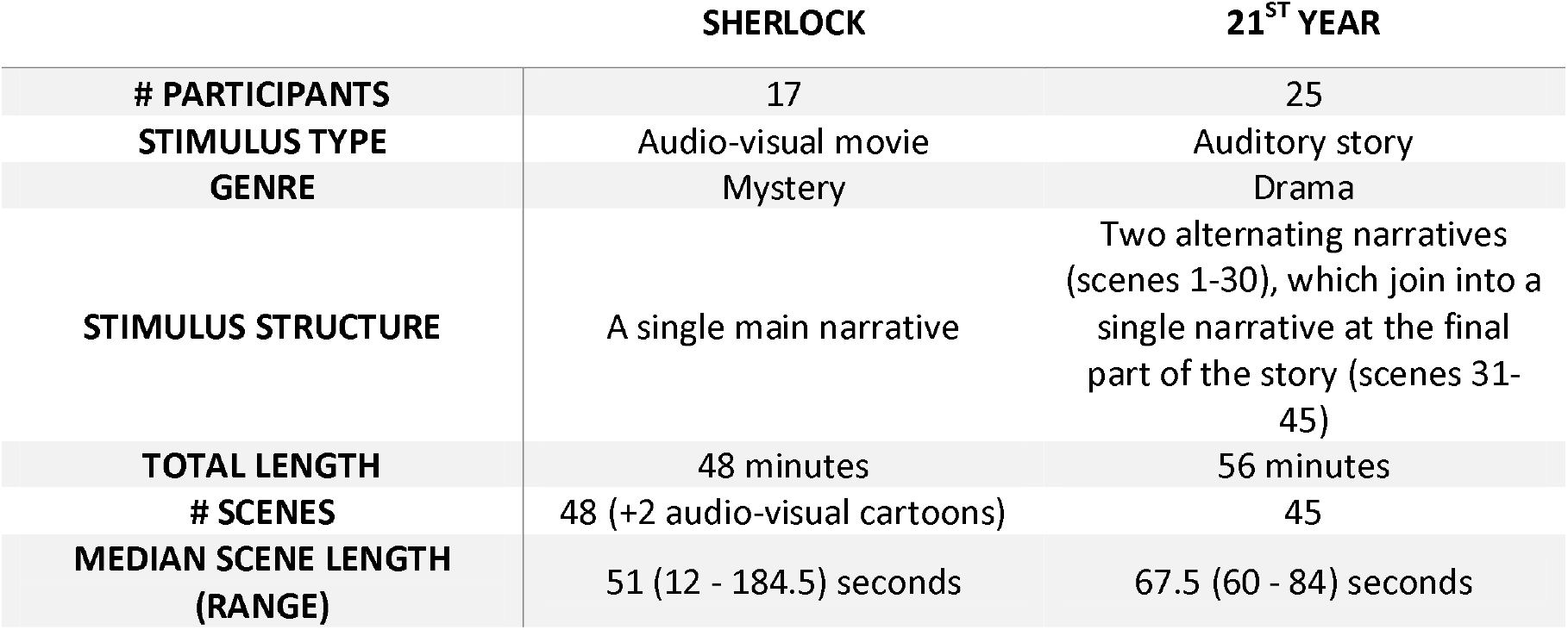
Details of datasets.

**Figure 1.**
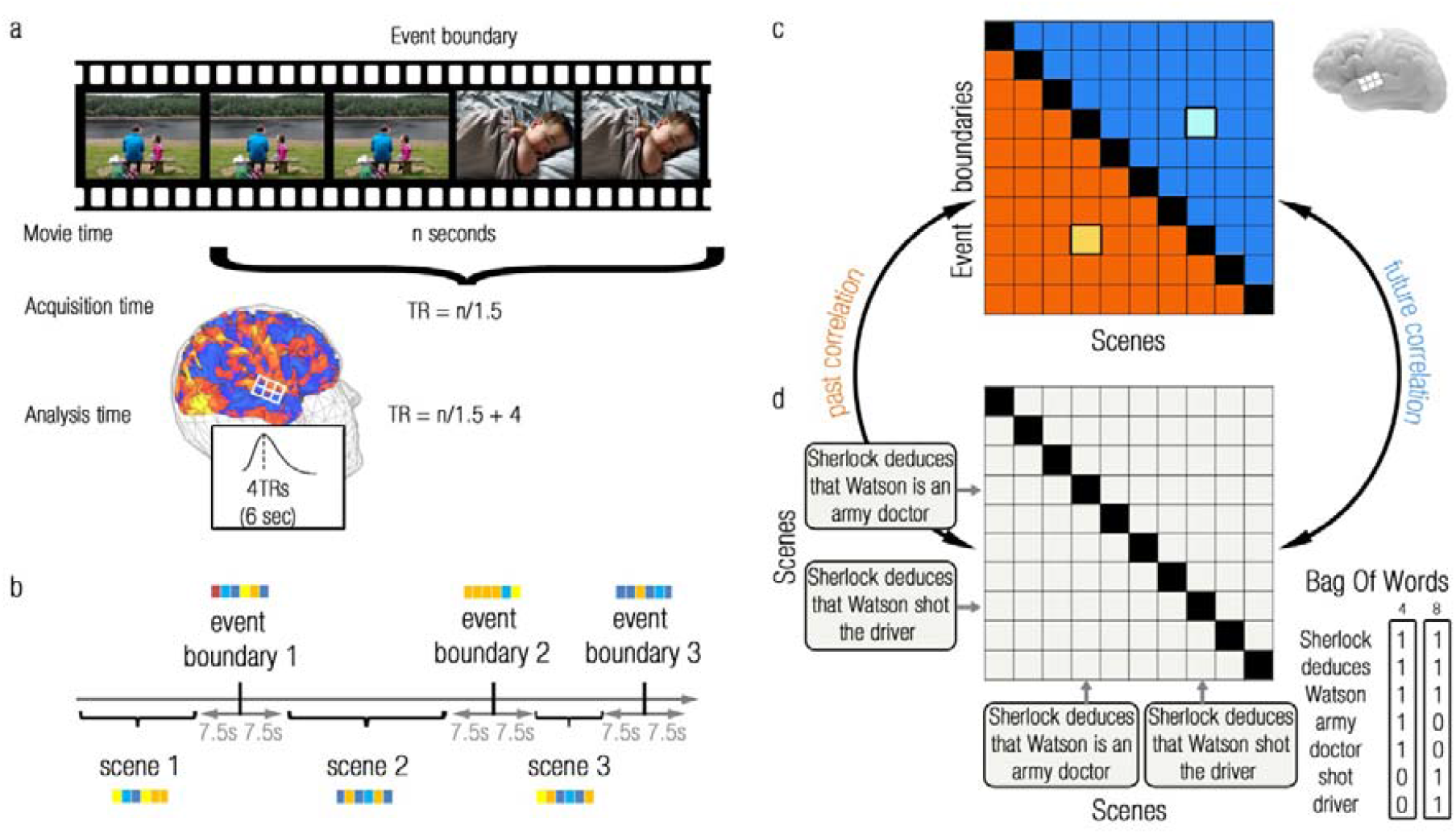
Schematic description of the analysis pipeline. **(a)** Participants watched a movie or listened to a story while being scanned. We defined event boundaries as the TRs in which scenes transitioned (TR=1.5 sec). We next shifted these TRs by 6 seconds (4 TRs) to account for the hemodynamic delay. **(b)** For each sphere in the brain (as illustrated in (a)), we extracted fMRI representations (patterns of voxel activity-levels) of event boundaries (black vertical lines). To separate between the BOLD signals of event boundaries and scenes, we defined the representations of scenes as the average representation across all within-scene TRs that are 5 TRs (7.5 seconds) remote from adjacent event boundaries. **(c)** We created a similarity matrix by cross-correlating the representations of event boundaries and scenes. The lower/upper triangular parts of the similarity matrix contained the similarities between the representations of event boundaries and the representations of their preceding/following scenes (past/future parts of the matrix, marked in orange/blue colours, respectively). We removed the main diagonal, which contained the similarity between each event boundary and its preceding scene, in order to detect only reactivation of temporally-remote scenes. Note that if similarities between event boundaries and scenes are a mere reflection of similarities between the scenes preceding the event boundaries and other scenes, this similarity would be reflected in both the past and future parts of the matrix. For example, if the similarity between event boundary 8 and scene 4 (entry (8,4) marked in bright orange) in fact reflected the similarity between scene 8 and scene 4, then the same similarity should be evident in the future part of the matrix (entry (4,8) marked in bright blue). To control for this possibility, we defined a reactivation index as the difference between the means of the past and future parts of the matrix, thus cancelling out any symmetric scene-similarity effects. **(d)** Bag Of Words analysis. We represented each scene according to the words it contains (illustrated for scenes 4 and 8). This representation was based on the occurrences of words, from across the entire narrative, that were used in each scene (illustrated in the bottom right for scenes 4 and 8). We then cross-correlated these scene-representations to create a symmetric context-similarity matrix, from which we removed the main diagonal. Next, we correlated the past/future part of the neural similarity matrix with the past/future part of the context similarity matrix, respectively. Finally, we subtracted the future correlation coefficient from the past correlation coefficient, thus cancelling out any effects resulting from scene-similarities. Note that we used some variations of this method for different datasets and analysis method, as described in the Online Methods.

See Figure 1 for a schematic of the analysis pipeline, and Online Methods for full methodological details, simulations and experimental constraints of our analytic methods.

### Reactivation of remote past events at event boundaries

We first tested for the reactivation of remote events within a single brain. To this end, we looked for brain regions where representations at event boundaries are more similar to representations of pasts scenes compared to future scenes (significantly positive reactivation index). Visual inspection of the results, as presented in Figure 2a, revealed that brain regions showing consistent effects across datasets included the bilateral precuneus/retrosplenial cortex (PCUN), angular gyrus/lateral Occipital Cortex (Ang/LOC) and posterior hippocampus (HIP). These regions were consistently observed in each of the two datasets and in a mega-analysis that pooled the participants of the two datasets together (The top two rows of Figure 4a displays the overlap between these regions across datasets. The statistical quantification of this replication will be presented in the section “Reactivation is specific to event boundaries” and Figure 4b/c). No regions showing the opposite effect, of a significantly negative reactivation index, were revealed. All maps have been cluster corrected for multiple comparisons across the entire brain (p<0.005, permutation test).

**Figure 2.**
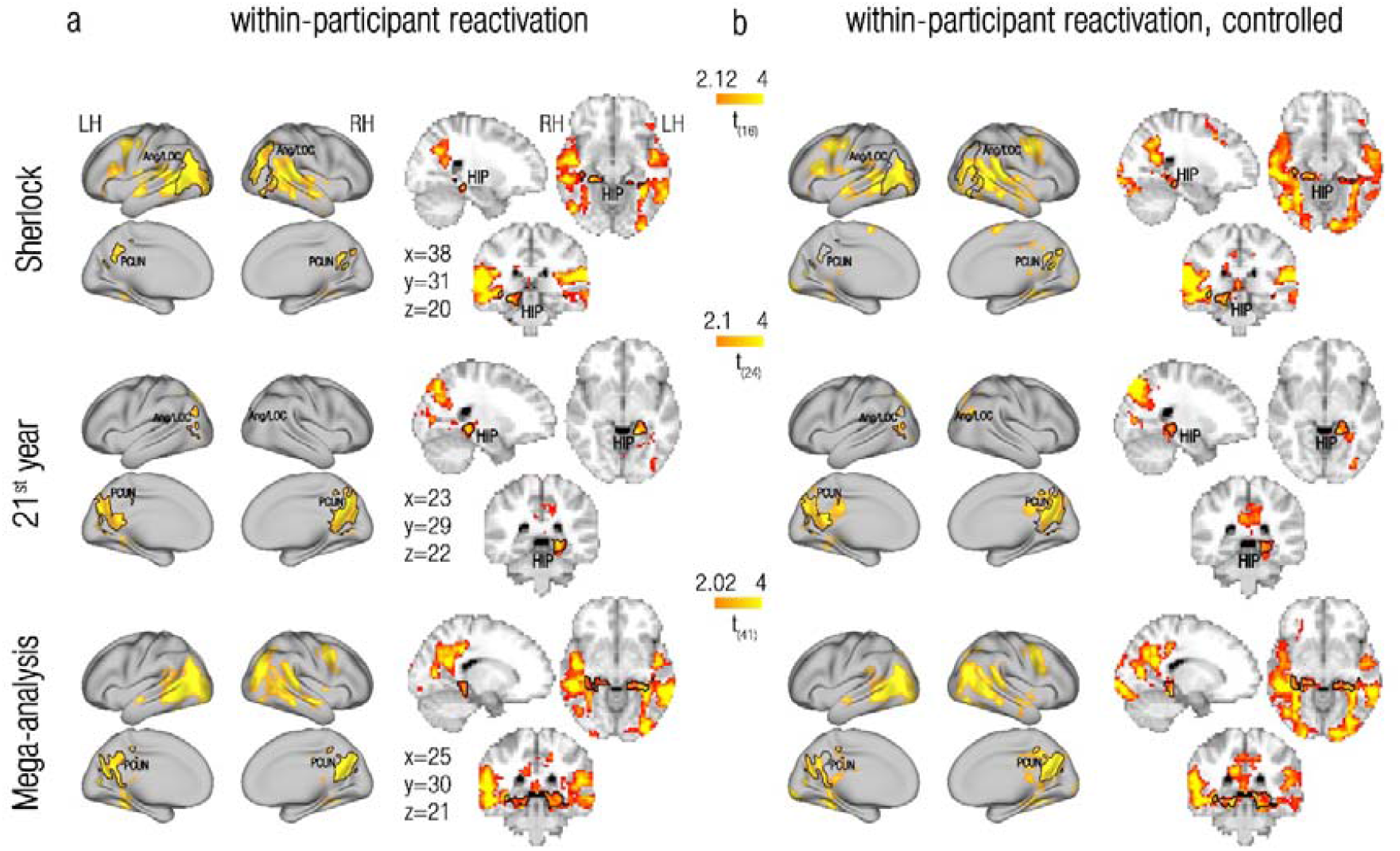
Within-participant reactivation of remote past events at event boundaries. **(a)** Within-participant reactivation, where reactivation indices were derived from event boundary and scene representations that were extracted from the same brain. **(b)** Controlled within-participant reactivation, where the within-participant reactivation indices at event boundaries were contrasted with those computed for control timepoints. Rows depict within-dataset group maps of the Sherlock dataset (1^st^ row), the 21^st^ year dataset (2^nd^ row) and a mega-analysis pooling together the two datasets (3^rd^ row). Maps are presented on both flat cortical surface and on 3D slices (dataset-specific MNI coordinates are depicted). Across all datasets, reactivation of temporally-remote past events (as reflected in positive t-values) was consistently found in the bilateral precuneus/retrosplenial cortex (PCUN), the Angular gyrus/Lateral Occipital Cortex (Ang/LOC) and hippocampus (HIP). To illustrate the resemblance in results between the uncontrolled and controlled analyses, we superimposed regions of interest based on the uncontrolled within-participants analyses of each dataset ((a), marked in black contours) on the corresponding controlled reactivation maps (b). LH, left hemisphere; RH, right hemisphere. All maps were cluster-corrected for multiple comparisons across the entire brain (p<0.005).

Note that, although regional variations between the group reactivation maps were also revealed, we refrain from interpreting results that do not show a qualitative replication across the 2 independent datasets. This is because such differences could be attributed to any of the design- or processing-related specifics of the individual datasets (see Table 1 and Online Methods).

### Reactivated representations have structure at a mesoscopic scale

Replay in rodents is typically studied at the single cell resolution. The human reactivation results presented so far were voxel-specific, which is the highest possible spatial resolution when using fMRI. However, naturalistic fMRI findings indicate that the neural representations of scenes are shared across participants^39-41^. Since there is no accurate correspondence between single voxels across brains (due to inter-brain anatomical/functional variability and image preprocessing constraints), it is likely that between-participant analyses do not sample the exact same voxels across brains, but rather voxels that are in some anatomical vicinity to each other. The findings of shared scene representations across brains therefore demonstrate that the representations of complex naturalistic events can be expressed at coarser scales, including mesoscopic scales, in which functional anatomy is consistent across participants. We were therefore interested to study whether reactivation is expressed at a mesoscopic resolution. Unlike in previous studies, we tested, not whether mere scene-representations are shared, but rather whether the reactivated representations observed at event boundaries within a single brain would also be shared across participants. To this end, scene representations extracted from one brain were correlated with the averaged event-boundary representation across the remaining participants of the same dataset.

Visual inspection of the results, as depicted in Figures 3a, demonstrates that the between-participant analysis indeed revealed reactivation in the bilateral precuneus/retrosplenial cortex, angular gyrus/lateral occipital cortex and bilateral posterior hippocampus, similar to the effects seen in the within-participant analysis (Figures 2a. The bottom two rows of Figure 4a displays the overlap between these regions across datasets. The statistical quantification of this replication will be presented in the section “Reactivation is specific to event boundaries” and Figure 4b/c). For the 21^st^ year dataset, these results were equally strong if only the congruent parts of the similarity matrix were analysed, but not if a Time (past/future) X Congruency (same/different narrative) interaction was used as a reactivation index (see sup. Figure 1 and Discussion). Similar results were also obtained for both datasets using the more typical scene-based between-participant analysis, in which the representations at event boundaries (rather than scene representations) extracted from one brain were correlated with the averaged scene (rather than event-boundary) representations across the remaining participants of the same dataset (Supplementary Figure 2, left column).

**Figure 3.**
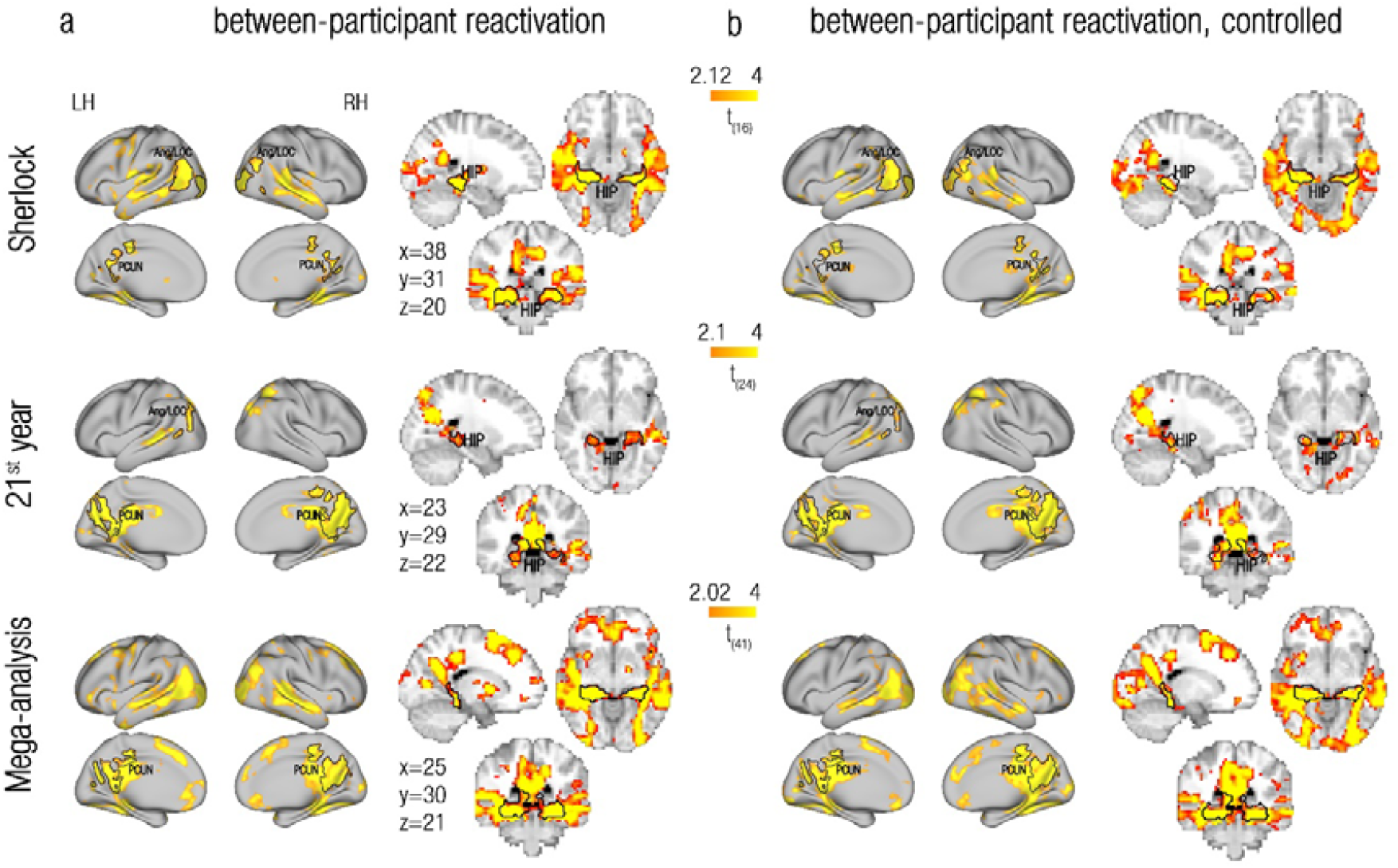
Between-participant reactivation of remote past events at event boundaries. **(a)** Between-participants reactivation, where reactivation indices were derived from the scene representations of one participant and the event boundary representations from all other participants of the same dataset. **(b)** Controlled between-participant reactivation, where the between-participant reactivation indices at event boundaries were contrasted with those computed for control timepoints. Rows depict within-dataset group maps of the Sherlock dataset (1^st^ row), the 21^st^ year dataset (2^nd^ row) and a mega-analysis pooling together the two datasets (3^rd^ row). Maps are presented on both flat cortical surface and on 3D slices (dataset-specific MNI coordinates are depicted). Across all datasets, reactivation of temporally-remote past events (as reflected in positive t-values) was consistently found in the bilateral precuneus/retrosplenial cortex (PCUN), the Angular gyrus/Lateral Occipital Cortex (Ang/LOC) and hippocampus (HIP). To illustrate the resemblance in results between the uncontrolled and controlled analyses, we superimposed regions of interest based on the uncontrolled between-participants analyses of each dataset ((a), marked in black contours) on the corresponding controlled reactivation maps (b). LH, left hemisphere; RH, right hemisphere. All maps were cluster-corrected for multiple comparisons across the entire brain (p<0.005).

We interpret the greater correlations between representations at event boundaries and past scenes (compared to future scenes) as reactivation of past information. However, a possible confound to this interpretation is a case where event representations are rapidly “flushed” after event boundaries^42^. If this is true, our effects may be due to representation at the event boundaries resembling the pre-flush representation more than the post-flush representation. This can be addressed by excluding, not only the pre-boundary event, but also the post-boundary event from the analysis (as already done in the within-participant analysis to control for autocorrelation-related biases). Doing so yields results that are highly similar to the original ones (sup. Figure 3), suggesting that flushing dynamics do not drive our effects.

Thus, our results so far reveal a set of brain regions in which remote scene-representations are reliably reactivated at the event boundaries of naturalistic fMRI data in humans. The reactivated representations are evident not only within a single brain, but they are also shared across brains, providing evidence that the reactivated representations have structure at the mesoscopic scale.

### Reactivation is specific to event boundaries

We have so far shown reactivation of past events at event boundaries, but is this effect specific to event boundaries? To test this, we repeated the reactivation analysis, but this time contrasted the reactivation indices at event boundaries with those calculated for control timepoints, that are adjacent to the event boundaries (see Online Methods and sup. Figure 4). As presented in Figure 2b and Figure 3b (and also see sup. Figure 2 for results of the scene-based between-participant analysis), this analysis revealed effects at a very similar set of regions compared to the original, uncontrolled reactivation analysis (Figure 2a, 3a, sup. Figure 2). Indeed, overlaying the ROIs defined from the uncontrolled analyses onto the maps resulting from the controlled analyses revealed a substantial overlap. This was true across datasets and analysis methods, attesting to a robust and specific effect of reactivation of remote past information to event boundaries.

**Figure 4.**
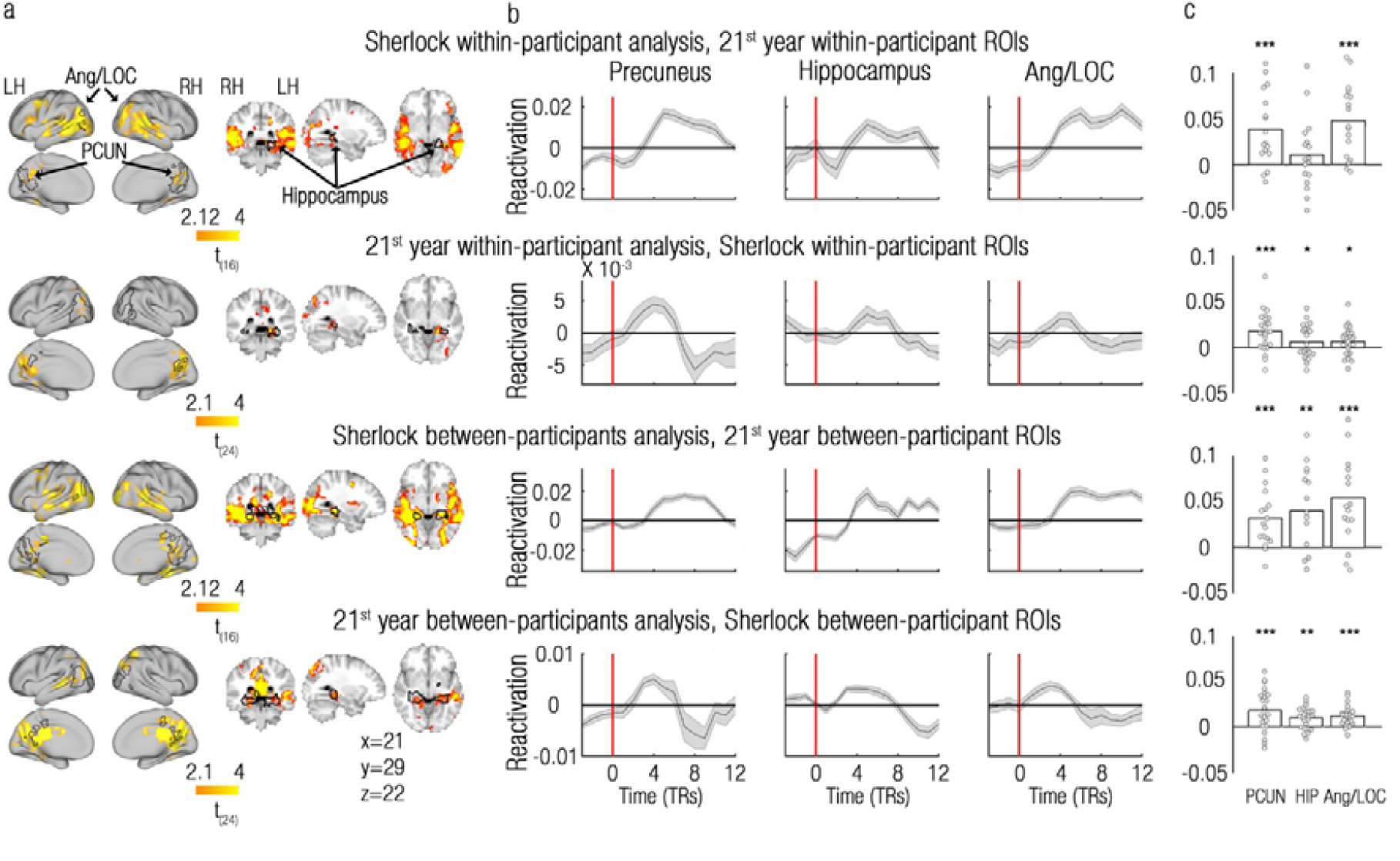
Reactivation is specific to event-boundaries – replication across datasets. **(a)** Group maps of each dataset (Sherlock/21^st^ year) superimposed with the ROIs defined from the other dataset, using the same analysis method (within/between-participant analysis). For example, in the first row, the within-participant reactivation group map of the Sherlock dataset (Figure 2a, 1^st^ row) is superimposed with ROIs defined from the within-participant reactivation group map of the 21^st^ year dataset (Figure 2a, 2^nd^ row). **(b)** Group-averaged reactivation indices derived from the ROIs in (a) are presented for the time period of 3 TRs prior to event boundaries until 12 TRs after event boundaries, uncorrected for the HRF delay. Event boundaries are depicted as red vertical lines. Grey shadings represent SEMs. (c) ROI analysis used to quantify the significance in (b). Reactivation indices are derived from each of the datasets and ROIs presented in (a) and TRs 1-6 (minus baseline) presented in (b). Single-participant values are represented as circles and group means are represented as bars. PCUN, Precuneus/retrosplenial cortex; Ang/LOC, Angular gyrus/Lateral Occipital Cortex; HIP, Hippocampus. *p<0.05; ** p<0.01; *** p<0.001.

The specificity of reactivation to event boundaries in the precuneus/retrosplenial cortex, posterior hippocampus and angular gyrus/lateral occipital cortex is clearly evident by examining the reactivation time-courses in these regions (the reactivation index at each scene-TR around the event boundary, see Online Methods), presented in Figure 4 (also see supplementary Figure 5 for results of the scene-based between-participant analysis). Across these brain regions, the reactivation time-courses peaked 6-9 seconds after event boundaries (vertical red lines), in a timecourse compatible with a BOLD response.

**Figure 5.**
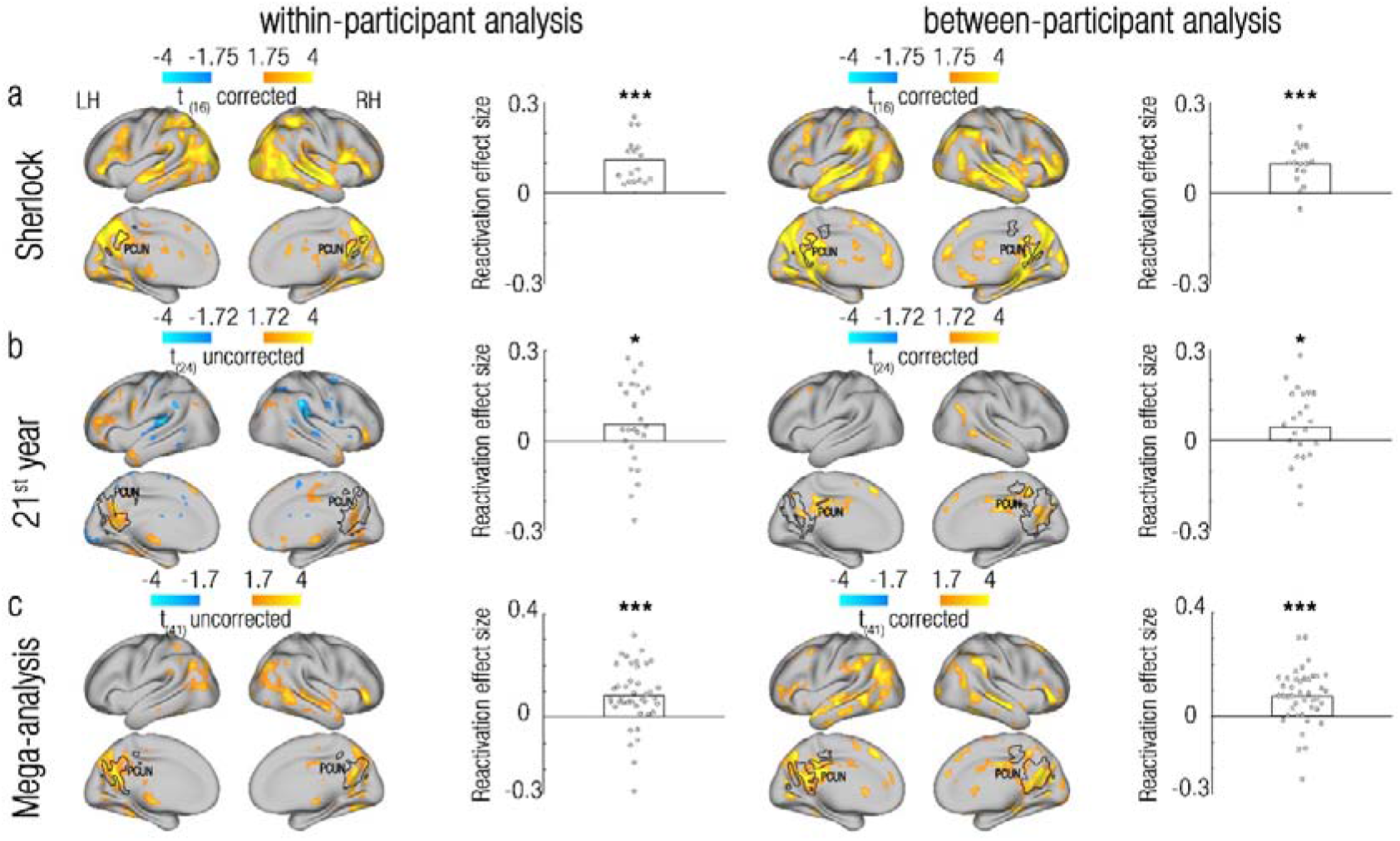
Relevant past events are preferentially reactivated at event boundaries. We measured the level to which reactivation is modulated by semantic context (as defined by a Bag Of Words model, Figure 1d) across the entire brain. Rows show within-dataset results of the Sherlock dataset **(a)**, the 21^st^ year dataset **(b)**, and a mega-analysis pooling together the two datasets **(c)**. Columns depict results obtained using different analysis methods (within/between-participant analysis). All maps were cluster-corrected for multiple comparisons across the entire brain (p<0.005), except for the within-participant analysis of the 21^st^ year dataset (b, 1^st^ column), and the within-participant mega-analysis (c, 1^st^ column). Maps are projected onto a flat cortical surface. Across datasets and analysis methods, the precuneus/retrosplenial cortex showed a consistent positive modulation of reactivation by semantic context (scenes with semantic context similar to that of each event boundary were reactivated more than scenes with different semantic contexts). To confirm that the same regions that showed a reactivation effect also showed a significant modulation of reactivation by semantic context, we tested independent precuneus/retrosplenial cortex ROIs (depicted in black contours), separately defined for each dataset (Sherlock/21^st^ year) and analysis method (within/between participant) in the reactivation analysis (Figure 2a, Figure 3a). We calculated the participant-specific reactivation effect sizes in these ROIs as the Time (pats-future) main effect / Time X Congruency (same/different story) interaction in the Sherlock/21^st^ year datasets, respectively (see Online Methods). Single-participant effect sizes are represented as circles and group means are represented as bars. LH, left hemisphere; RH, right hemisphere; * p<0.05; *** p<0.001.

Of importance, quantifying the effects observed in these ROI-specific reactivation timecourses also allowed us to test statistically whether the same brain regions are indeed detected across the two independent replications of our results. We conducted a direct cross-validation across datasets (Sherlock/21^st^ year) by defining ROIs in one dataset and testing the magnitude of their reactivation signals in the other dataset, using the same analysis method (Figure 4). The tested brain regions showed highly replicable effects across these 2 independent datasets, using either analysis method, as reflected in the test-specific p-values (supplementary Table 1, the only exception is the hippocampus in the within-participant analysis of the Sherlock dataset, p=0.1). Meta-analysis of p-values across datasets showed significant results for each and every brain region (within participant analysis: precuneus/retrosplenial cortex: χ^2^_(4)_=27.63, p<0.001; posterior hippocampus: χ^2^_(4)_=10.14, p=0.038; Ang/LOC: χ^2^_(4)_=21.36; p<0.001, between participant analysis: precuneus/retrosplenial cortex: χ ^2^_(4)_=27.63, p<0.001; posterior hippocampus: χ2_(4)_=23.47, p<0.001; Ang/LOC: χ2_(4)_=27.63; p<0.001, Fisher’s test, α=0.017 for all tests). Very similar results were obtained in a cross-validation analysis across datasets using the scene-based between-participant analysis (precuneus/retrosplenial cortex: χ^2^_(4)_=27.63, p<0.001; posterior hippocampus: χ^2^_(4)_=23.47, p<0.001; Ang/LOC: χ^2^_(4)_=25.43, p<0.001, Fisher’s test, α=0.017 for all tests; supplementary Table 1, supplementary Figure 5). The reactivation effect is therefore not only specific to event boundaries, but also highly replicable across datasets.

### Relevant past events are preferentially reactivated at event boundaries

Our results so far indicate a robust and reliable reactivation of remote past events, which is specific to event boundaries of ongoing naturalistic experience. But is there also specificity to the content of reactivated information? Is *relevant* past information reactivated more than irrelevant past information? To test this hypothesis, we constructed a Bag Of Words model of the text in each narrative (the scene descriptions and dialogues of the Sherlock movie and the full 21^st^ year story). Each scene was represented by a vector, whose elements represented the number of occurrences of each word. For example, the Bag-Of-Words vector for this paragraph would have a 3 in the element that represents the word “vector” (see illustration at the bottom right of Figure 1d). Under the assumption that similar contexts share similar words, by correlating these Bag Of Words representations across scenes, we created a context similarity matrix for each experimental stimulus (Sherlock movie description/21^st^ year story, see illustration in Figure 1d, and similarity matrices in sup. Figure 6b/d). Next, for each sphere in each participant’s brain, we correlated the neural event-boundary X scene similarity matrix with the dataset-specific Bag Of Words scene-similarity matrix: for the Sherlock dataset, we correlated the past and future parts of the matrices separately and calculated a time (past-future) main effect. For the 21^st^ year dataset, which contained two alternating narratives, we computed correlations between the past congruent (same narrative), past incongruent (different narrative), future congruent and future incongruent entries of the matrices, and computed a time X congruency interaction (see Online Methods).

In the Sherlock dataset, a wide network of brain regions (including the precuneus/retrosplenial cortex, angular gyrus/lateral occipital cortex and hippocampus) showed a positive modulation of fMRI reactivation by context-similarity across analysis methods (Figure 5a). The 21^st^ year dataset showed weaker effects (as expected due to methodological constraints, see Online Methods), which replicated across datasets only in the precuneus/retrosplenial cortex (whole-brain corrected in the between-participant, but not the within-participant analysis, Figure 5b). This effect in the precuneus/retrosplenial cortex was also evident in a mega-analysis across datasets (whole-brain corrected in the between-participant, but not the within-participant analysis, Figure 5c. For additional results on the entire context-similarity matrices, see Online Methods and sup. Figure 6). Significant results were also obtained using the scene-based between-participant analysis (sup. Figure 7). These results suggest that in the precuneus/retrosplenial cortex, reactivation at event boundaries is selective for remote past events that are relevant (have similar context) to the current situation.

Whilst it is encouraging that most of these effects survive whole-brain correction, the critical question is whether the same voxels that reactivate past scenes on average (Figure 2, Figure 3), also do so selectively. To test this, we used an independent ROI analysis. We defined precuneus/retrosplenial cortex ROIs, separately from each dataset (Sherlock/21^st^ year) and analysis method (within/between participant) in the reactivation analysis (Figure 2a, Figure 3a) and tested the context specificity (Figure 5, sup. Figure 7) within this ROI in the same dataset and using the same analysis method. Precuneus/retrosplenial cortex ROIs showed significant context-specific reactivations with each approach and dataset (Sherlock within participant analysis: t=5.98, p<0.001, Sherlock boundary-based between participant analysis: t=4.86, p<0.001, Sherlock scene-based between participant analysis: t=10.55, p<0.001, 21^st^ year within participant analysis: t=1.92, p=0.02, 21^st^ year boundary-based between participant analysis: t=1.51, p=0.039, 21^st^ year scene-based between participant analysis: t=2.11, p=0.019, one-tailed permutation tests, see sup. Table 2 for means, SEMs and effect sizes). A meta-analysis conducted across the two independent datasets for each analysis method confirmed a robust replication of this effect (within participant analysis: χ^2^(4)=21.64, p<0.001; boundary-based between participant analysis: χ^2^(4)=20.3, p=0.001; scene-based between participant analysis: χ^2^(4)=21.36, p<0.001, Fisher’s test). Mega-analyses, pooling together all participants from the two datasets also confirmed these results (all p<0.001, Figure 5c, supplementary Figure 7c). Thus, the reactivation seen in the precuneus/retrosplenial cortex at event boundaries (Figure 2, Figure 3) is selective for relevant past information.

## Discussion

How do we make sense of our ongoing experiences? Our findings propose that past information that is relevant for the comprehension of the current situation is reactivated at event boundaries. We found evidence for reactivation of past events in the same brain regions (precuneus/retrosplenial cortex, posterior hippocampus, angular gyrus/lateral occipital cortex) across 2 independent datasets, that are vastly different from one another (Table 1), suggesting that this reactivation represents a general attribute of the processing of ongoing experiences. These results, observed both within and between individuals, demonstrates that this reactivation exists both at the local (voxel-specific) scale and at the mesoscale (Figures 2 and 3). We further established that the observed reactivation is not only specific to event boundaries (Figure 4), but also to selected past events: the reactivation of events in the precuneus/retrosplenial cortex is modulated by semantic context, such that past events with a similar context as the current event were more likely to be reactivated (Figure 5). Taken together, these findings demonstrate that reactivation of past events at event boundaries is a selective mechanism that can piece together relevant parts of an ongoing experience.

Previous studies of replay have used experimental designs based on multiple repetitions of the same task^13, 14, 43^. Such experiments demonstrated that past knowledge (e.g. the pre-learned structure of a maze) and current learning (e.g. the position of a new reward location in the maze) can be combined through replay^32, 43^. However, the documented replay was not selective for specific past events, but rather represented a whole knowledge-structure, acquired across repeated trials. In other words, replay has been shown to reflect the assimilation of information into a previously-acquired schema. Contrary to such over-trained experiments, in the naturalistic processing of a narrative, each event is only experienced once. Accordingly, we designed an analysis method which allowed us to treat events as unique pieces of information whose inter-relations can be inferred as a narrative unfolds. Indeed, our observation of a selective reactivation of unique events that are semantically-related to one another suggests the formation of relations between these individual events. Inferring these relations lies at the heart of understanding unfolding events, and we therefore suggest reactivation as the mechanism that constructs this relational understanding while events unfold.

Reactivation was observed in the hippocampus and in regions of the DMN. Indeed, replay and sharp wave ripples have been documented simultaneously in these brain regions in the human^26, 27^, primate^28^ and rodent brain^25, 29^. Interestingly, replay/reactivation are commonly interpreted as supporting memory of past events and planning of future behavior^13, 30^. The DMN is also commonly believed to be an “offline” (resting-state) network, involved in remembering the past and envisioning the future, but to be disengaged from processing immediate external information^44, 45^. However, in naturalistic experiments, the DMN has been hypothesized to have an online function in interpreting ongoing information in light of prior knowledge^9, 10, 39, 46^, though the mechanism underlying the continuous binding of information has not yet been characterized. The naturalistic paradigm used here in combination with the mechanistic insights offered by our analytic approach indicate that, during an unfolding experience, the DMN processes *online* information by employing a reactivation mechanism that binds the present with related past events. The online nature of this computation can therefore construct our understanding of current events. Note that, it is not under dispute that reactivation or the DMN support offline functions, such as memory or planning. The online-constructed links between events could build our understanding of the present, and simultaneously be consolidated into memory and inform subsequent decisions. We therefore suggest that reactivation in the DMN may do more than consolidating *past* events into memory or informing *future* decisions; it may serve as an information-binding mechanism that underlies the understanding of the *present*.

Not all regions of the DMN have been detected to reactivate remote past information in our results. However, both implicated nodes, the precuneus/retrosplenial cortex and the angular gyrus, have been highlighted as particularly likely to represent event models^4, 5, 47^. In addition, these two regions have been shown to have long timescales of neural dynamics^9, 47, 48^, and were therefore suggested to be able to intrinsically retain past information throughout an uninterrupted movie viewing^9^. It is therefore possible that, at event boundaries, the precuneus/retrosplenial cortex and the angular gyrus can reactivate specific remote event-models out of all past models they hold for the duration of the movie. Interestingly, the detection of neural signatures of event boundaries in these particular regions has been demonstrated to predict hippocampal response during movie watching^47^. It is therefore possible that, after the selection of events to be reactivated in these DMN regions, the hippocampus can bind these events with the current event-model, resulting in an inter-connected representation of an unfolding experience^7, 49-52^.

These regions were reliably detected in both datasets using a time main effect as a reactivation index, but not when using a time X congruency interaction in the 21^st^ year dataset. This could be due to the fact that the 21^st^ year story has an alternating narrative structure. At the end of each narrative-specific scene, the listeners likely predict that the following scene will be of the other (incongruent) narrative. It is therefore plausible that event boundaries will not only trigger reactivations that encode the content of the just-ended scene, embedded in its narrative. Event boundaries may also trigger reactivations of the incongruent narrative, to set the context for the predicted upcoming scene. If this is true, event boundaries would display reactivations of both narratives (congruent and incongruent with the just-ended scene), that will be evident in the time main effect but not in the time X congruency interaction. Future studies will be needed to investigate interaction effects in a story that contains two narratives with an unpredictable structure.

We note that, of the three brain regions detected to reactivate past information, only the precuneus/retrosplenial cortex showed a context-selective modulation of this reactivation which was reproducible across datasets. The retrosplenial cortex is indeed believed to process relational and contextual information that is needed for the consolidation of events into memory^53^. In addition, naturalistic experiments have repeatedly identified the precuneus/retrosplenial cortex as the main brain region involved in context processing^9, 39, 46^ and in the representation of semantically-central events (showing dense semantic relations to other events) during recall of a narrative^54^. It therefore seems reasonable that, as our results indicate, the precuneus/retrosplenial cortex has a unique contribution to the selection of contextually-relevant events for reactivation during narrative processing. Nevertheless, our results also hint that other brain regions may take part in this process, as evidenced by the results obtained from the Sherlock dataset (Figure 5a). Since these other brain regions were not replicated in the 21^st^ year dataset, future work, using a larger number of datasets, will be needed to target the contextual modulation of reactivation in additional brain region.

It is also noteworthy that our analyses detected regions that reactivated past events, but no regions where representations at event boundaries were more similar to (and hence predictive of) future events. In theory, such predictive coding could have been detected in our analyses, since prediction and planning for the near future have been postulated to be integral to perception, and to be represented in event models^1, 2, 5^. Indeed, in the rodent literature, replay has been documented to inform immediate goal-directed choices^32, 55, 56^ (but also see^57^). Future studies will be needed to characterize the conditions under which predictive coding could also be detected in naturalistic narrative processing experiments in humans.

In conclusion, by the end of this paper, the reader’s brain should have constructed an interconnected set of representations, relating our experimental hypotheses (presented in the Introduction), our findings (presented in the Results), and our interpretations of these findings, as just described (we assume not all readers will have read the Methods section). We propose that hippocampus and DMN regions in the reader’s brain formed this complex structure of knowledge via a selective reactivation of relevant information at event boundaries, such as endings of paragraphs. If this knowledge structure has indeed been formed during the reading, the reader should have been able make sense of this paper, and perhaps even remember it in the long run, or make use of it in her/his own future work.

## Acknowledgements

We would like to thank Marlene Behrmann, Aya Ben-Yakov and Tamar Makin for their valuable insights, which greatly contributed to the manuscript. AH would like to thanks baby Adam, for allowing his mum to work on this paper during her maternity leave.

## Online Methods

### Overview

Here, we aimed to study reactivation of temporally-remote events (i.e. further than the just-ended event) at event boundaries of naturalistic stimuli. For this purpose, we analyzed two naturalistic datasets, where participants either watched a movie or listened to a story while undergoing a whole-brain fMRI scan. We measured reactivation as the correlation between fMRI representations at event boundaries (the endings of scenes, Figure 1a), and representations of previous scenes (Figure 1b). The lower/upper triangular parts of the resulting similarity matrix reflected similarities between each event boundary and earlier/later scenes (Figure 1c, orange/blue, respectively).

We wanted to look for reactivation of remote scenes, so we removed the immediately preceding scene of each event boundary. However, one possibility is that each event boundary correlates with its immediately preceding scene, and the preceding scene correlates with other scenes. Thus, correlations between event boundaries and remote scenes may in fact reflect between-scene correlations, rather than reactivation of remote information. Yet, correlating between scenes results in a symmetrical similarity matrix, in which the same inter-scene correlations exist for both past scenes (scene 8 vs. scene 4) or future scenes (scene 4 vs. scene 8, Figure 1c). It is therefore possible to control perfectly for this potential confound by computing a reactivation measure that *contrasts the past with the future* (thus cancelling out symmetric similarities). By computing this reactivation index using a whole-brain searchlight approach (namely, using a sphere centred on each fMRI voxel in each participant’s data), we were able to test in which brain regions representations at event boundaries correlate with (i.e. reactivate) past scene-representations.

In this within-participant analysis, we calculated correlations between representations at event boundaries and scenes, extracted from a single brain. We also performed a between-participant analysis, in which representations of scenes were extracted from the brain of one participant and correlated with the averaged event-boundaries representations extracted from the brains of the remaining participants of the same dataset. Note that for the within-participant reactivation analysis we needed to carefully control for autocorrelations in the raw data, since representations of event boundaries and scenes were extracted from the same fMRI signals. This problem will be discussed in the section “Autocorrelations in within-participant analyses”, and does not apply to the between-participant analysis, which correlates representations between signals of independent participants.

Finally, to test whether reactivation is selective for scenes with relevant semantic context, we used the overlap in words between scenes to further characterize the observed reactivation (Bag Of Words model, Figure 1d).

### Data

This study reports findings from two naturalistic fMRI datasets (see Table 1): The Sherlock dataset^40^ downloaded from Princeton University’s DataSpace repository^37^; and the 21^st^ year dataset^39^, downloaded from the Narratives repository^38^.

#### Participants

*Sherlock*. 22 participants (10 female, age range 18-26) were recruited for the original study, of which 17 met the original study’s inclusion criteria, and were therefore also analysed in the current study.

*21^st^ year*. 25 participants (14 female, age range 18-40) were analysed in the original and current study.

All participants were right handed native English speakers, and none had been exposed to the experimental stimulus prior to the scanning session.

#### Experimental stimuli

##### Sherlock

Here we analysed data from participants who watched the first 48 minutes of Episode 1 of the BBC’s television series “Sherlock”. The plot had a continuous structure, with varying scene-lengths (median 51 seconds, range 12 - 184.5 seconds). Scene timings were taken from the original study, where they were manually defined based on major shifts in the narrative^40^. The experimental stimulus was divided into two segments of 23/25 minutes, and a 30 seconds audiovisual cartoon (unrelated to the main stimulus) was presented prior to each segment.

##### 21st year

Participants listened to a 56 minute-long story, which was segmented into scenes by design. These scenes had relatively constant lengths (median 67.5 seconds, range 60 - 84 seconds), and were separated by 4.5-6 seconds of silence. The narrative was composed of two unrelated storylines that alternated during the first 30 story-scenes, and then merged into a unified storyline during the last 15 story-scenes.

#### fMRI acquisition

##### Sherlock

Imaging was performed on a 3T full-body scanner (Siemens Skyra) with a 20-channel head coil. Functional images were acquired using a T2*-weighted echo planar imaging (EPI) pulse sequence (TR 1500 ms, TE 28 ms, flip angle 64°), each volume comprising 27 slices of 4 mm thickness, in-plane resolution 3 × 3 mm^2^, FOV 192×192 mm^2^), with ascending interleaved acquisition.

##### 21st year

Imaging was performed on a 3T full-body MRI scanner (Skyra, Siemens) with a 20-channel head coil. Functional images were acquired using the same parameters as in the Sherlock dataset. Anatomical images were acquired using a T1-weighted magnetization-prepared rapid-acquisition gradient echo (MPRAGE) pulse sequence (TR 2300 ms; TE 3.08 ms; flip angle 9°; 0.86 x 0.86 x 0.9 mm^3^ resolution; FOV, 220 x 220 mm^2^).

#### Data preprocessing

##### Sherlock

The fully preprocessed images from the Sherlock dataset were downloaded and used. Preprocessing was performed in FSL (http://fsl.fmrib.ox.ac.uk/fsl), and included slice time correction, motion correction, linear detrending, temporal high-pass filtering (140s cutoff), and coregistration and affine transformation of the functional images to an MNI template brain. Functional images were resampled to 3 mm isotropic voxels.

##### 21st year

Raw images from the narrative dataset were downloaded and then preprocessed using FSL and in-house Matlab code (version 2018a, Mathworks, Natick, MA, USA). Functional data were analysed using FMRIB’s expert analysis tool (FEAT, version 6). The following preprocessing steps were applied to each participant’s data: motion correction using FMRIB’s Linear Image Registration Tool (MCFLIRT), brain extraction using BET and high pass temporal filtering (120s cutoff). Functional images were aligned to structural images initially using FMRIB’s Linear Image Registration Tool (FLIRT)^58, 59^ and then optimized using Boundary-Based Registration (BBR)^60^. Structural images were transformed into MNI space using FMRIB’s Nonlinear Image Registration Tool (FNIRT) and the resulting warp fields were applied to the functional images. Tissue-type segmentation was carried out using FAST to create white matter/CSF nuisance masks. To avoid the inclusion of grey matter voxels in these nuisance masks, these masks included only voxels identified as white matter/CSF with probability of 1, restricted by an anatomically-based bounding box (-44<x<42, -84<y<42, - 4<z<34 for white matter; -42<x<38, -64<y<38, -22<z<28 for CSF), and eroded by a sphere of radius 5 and 2 mm for white-matter and CSF, respectively^61^. White-matter and CSF time-courses were extracted for each functional scan, and their contribution to the BOLD signal, as well as the contribution of motion parameters, was later removed. Functional images were resampled to 3 mm isotropic voxels.

Across all analyses, unless otherwise specified, fMRI BOLD responses were shifted by 4 TRs (6 seconds) in relation to the experimental stimuli to account for the canonical HRF delay (Figure 1a).

### Data analysis

#### fMRI representations

To test reactivation across the entire brain, a searchlight approach was employed, with spheres of 3 voxel (9mm) radius confined by a standard MNI mask. Extracting representations (time-specific voxel patterns) from each sphere necessitated the separation of fMRI BOLD signals of event boundaries from the signals of whole scenes. Therefore, as illustrated in Figure 1b, in each participant, representations of event boundaries were defined as the voxel pattern in the single TR that ends each scene. Representations of scenes were defined as the average representation across all within-scene TRs that are 5 TRs (7.5 seconds) remote from adjacent event boundaries. The selection of a 5 TR gap was based on the estimation that the canonical HRF peaks at 4-6 seconds (3-4 TRs) of stimulus onset, hence a minimal gap was chosen as to avoid the HRF peak but also to allow sufficient scene-TRs to be analysed.

Note that within-scene averaging of representations is a common practice^39-41, 62, 63^, which is consistent with the pattern of whole-scene reactivation at event boundaries seen in EEG data^17^. This averaged scene representation ignores rapidly changing scene-information and likely reflects more stable elements of the scene, or its gist, as believed to be captured by event models^4^.

Due to the need to separate between the BOLD signals of event boundaries and scenes, 5 scenes from the Sherlock dataset that were shorter than 10 TRs were excluded from further analyses, along with their corresponding event boundaries. In the same dataset, the two scenes corresponding to the unrelated cartoon added to the experimental stimulus were also removed, along with the final scene of each scan, which did not have enough TRs following it to account for the HRF delay.

#### Within/between-participant similarity matrices

In order to detect similarities in representations that would indicate information reactivation, the Pearson correlations between event boundary representations and scenes representations were calculated (Figure 1c). This yielded an event boundary X scene similarity matrix, in which the lower triangular part reflected correlations between each event boundary and its previous scenes, and the upper triangular part reflected correlations between each event boundary and its following scenes. The main diagonal of this matrix was removed, as it reflected similarities between each event boundary and its immediately preceding scene, rather than reflecting similarities with remote scenes.

Importantly, this similarity matrix was computed using two related analytic approaches: a within-participant analysis and a between-participant analysis. In the within-participant analysis, for each sphere in each participant, representations of event boundaries and of whole scenes were derived from a single brain, and correlated to create a within-participant (event boundary X scene) similarity matrix. Thus, this approach allowed the detection of reactivation of remote scene representations at event boundaries within a single brain.

However, under the assumption that brain activity is modulated by naturalistic stimuli in a similar fashion across participants, correlations between brains could also be calculated (inter-subject correlations^64^). This approach has been implemented in the analysis of naturalistic data, demonstrating that representations of scenes have structure at a meso-scale in which functional anatomy is shared across participants^39-41^. Hence, reactivation too may have a structure at a meso-scale, and could be detected in the similarity between the representations of event boundaries in one brain and the (shared) representations of scenes averaged across all other brains.

However, it is also possible that the representations at event boundaries may be shared across brains. This previously unexplored possibility would suggest that the reactivated pattern itself, rather than the representations of scenes, is common across all participants. To test this hypothesis, for each sphere in each participant, representations of scenes were extracted and correlated with the same-sphere event-boundary representations, which were averaged across all other participants of the same dataset. This resulted in an event boundary (group-averaged) X scene (of one participant) similarity matrix.

Note that, the brain regions showing reactivation would not necessarily be similar across these two types of analysis. First, the within-participant analysis should be equally sensitive for the detection of reactivation across all brain regions. This is unlike the between-participant reactivation, which could only be detected in brain regions in which representations are shared across participants. Second, compared to the within-participant analysis, the higher SNR of the between-participants analysis, resulting from the averaging of signals across many brains, may allow an easier detection of areas that show a reactivation effect (thus resulting in stronger effects).

Furthermore, autocorrelations needed to be controlled for in the within-participant reactivation analysis, since representations of event boundaries and scenes were extracted from the same fMRI signals (as will be discussed in the “Autocorrelations in within-participant analyses” section). However, in the between-participant analysis, the representations of event boundaries were temporally independent of the scene representations (as they came from different brains), and therefore no autocorrelation issues existed.

All following analyses reported here were conducted using both the within-participant approach and the boundary-based between-participant approach. In addition, results from the scene-based between-participant analysis, using an event boundary (of one participant) X scene (averaged across all other participants) similarity matrix, are presented in the supplementary materials.

Finally, it is possible that greater correlations between representations at event boundaries and past scenes (compared to future scenes) representations reflect event representations that are rapidly “flushed” after event boundaries^42^. This would entail that representations at the event boundaries resemble the pre-flush representation more than the post-flush representation. To rule out this possibility, the between-participant analyses were repeated while excluding the post-boundary events (in addition to the pre-boundary events). Note that this control analysis was not needed for the within-participant analysis, as the related events were already removed to control for autocorrelation-related biases (see section “Autocorrelation in within-participant analyses”).

#### Reactivation analysis

Correlations between event boundary- and scene-representations were meant to detect reactivation of remote scenes at event boundaries. However, it is also possible that these correlations would simply reflect similarities between scene representations, for two reasons: 1) due to the low temporal resolution of fMRI data, event boundaries are likely correlated with their surrounding scenes; 2) If each event boundary representation contains reactivation of its immediately preceding scene^17^, the representation at the boundary would be similar to the scene representation. Since same-narrative scenes are likely to have shared features, and therefore similar representations, the representation at the event boundary of a certain scene may be similar to remote scenes, not due to reactivation, but merely due to between-scene similarities (the event boundary representation is similar to its containing scene representation, the containing scene representation is similar to other scene representations). However, between-scenes similarities necessarily yield a symmetric similarity matrix, in which the same inter-scene correlations exist for both past scenes (e.g. scene 8 vs. scene 4) or future scenes (e.g. scene 4 vs. scene 8, Figure 1c). Therefore, to rule out the possibility that between-scene similarities were measured, rather than reactivation of remote scenes at event boundaries, a “reactivation index” was used. This index was defined as the difference between the mean of the lower triangular part of the similarity matrix (past part), and the mean of the upper triangular part of the similarity matrix (future part):

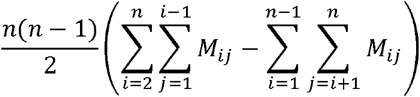

Where M is an n X n matrix, i and j are row and column indices, respectively.

This index controls for any effects of scene similarities, since it cancels out symmetric entries in the lower and upper triangular parts of the matrix, and thus isolates the asymmetric components of the matrix as a probe for reactivation.

For the 21^st^ year datasets, we also repeated the between-participant reactivation analyses using only the congruent entries of the similarity matrix, and also using a reactivation index based on a Time (past/future) X Congruency (same/different narrative) interaction.

Thus, our analytic approach consisted of the following steps: for each sphere in each participant, event boundary and scene representations were extracted. An event boundary X scene similarity matrix was calculated, from which the main diagonal was removed. Correlation coefficients in the matrix were Fisher transformed, and a reactivation index was then calculated, and assigned to the anatomical coordinates of the sphere center. This procedure was repeated for all brain spheres, creating a single-subject map. Within each of the Sherlock/21^st^ year datasets, single-subject maps were submitted to a voxel-wise two-tailed group t-test, creating a dataset-specific group map, thresholded at p<0.05. These group maps were corrected for multiple comparisons using a permutation test, using the null distribution of the maximal cluster mass, as implemented in FSL’s randomise, thresholded at p<0.005. Finally, the corrected maps were projected onto a template MNI brain using the connectome workbench.

#### ROI definition

Regions that showed consistent effects across datasets were detected visually. The 4 reactivation group maps (resulting from the Sherlock/21^st^ year datasets in the within/between participant analyses) were masked by relevant regions from anatomical atlases to define 4 sets of ROIs, as detailed below.

All ROIs were defined based on group maps in a statistically significant threshold, and after correction for multiple comparisons. Anatomical definitions of the precuneus and lateral occipital cortex (joining the superior and inferior sub-divisions) from Harvard-Oxford cortical structural atlas were used to mask the group maps, creating dataset/analysis-specific precuneus/retrosplenial cortex and angular gyrus/lateral occipital cortex ROIs. All hippocampal subregions of the Julich histological atlas were merged to create an anatomical definition of the hippocampus, which was used to mask all group maps and create dataset/analysis-specific hippocampal ROIs.

#### Time-specificity of Reactivation

To test whether the observed reactivation is specific to event boundaries, representations at event boundaries must be contrasted with those of a control time-point. But which control time point should be chosen? Event boundaries are by definition positioned between two adjacent scenes. A control time-point that precedes an event boundary will be much closer to the past period (as defined by the boundary) and a control time-point that follows an event boundary will be much closer to the future period. Due to autocorrelations in fMRI signals, the choice of each of these time-points to serve as a control will bias our analyses (as detailed in the section “Autocorrelations in within-participant analyses”). This problem can be tackled by choosing both a past and a future control timepoint that are both equally distant in time from the event boundary, and thus have symmetric time-biases that will cancel out.

Therefore, a balanced control was defined as the averaged reactivation indices computed for two symmetrical timepoints on both sides of each event boundary. Contrasting the reactivation indices of event boundaries with those of the balanced controls was defined as:

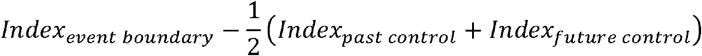

Control timepoints should be chosen such that they would not include any contribution from their adjacent event boundaries, otherwise the experimental effects seen at the event boundaries would be diminished by the contrast. Given the sluggish nature of the BOLD response, control points that are 10 TRs (15 seconds) distant from the event boundaries were chosen for the Sherlock dataset. This optimal choice, however, could not be made for the 21^st^ year dataset, given its experimental design. Since most event boundaries are preceded by a scene of the same narrative and followed by a scene from a different narrative, the past and future controls would potentially be contaminated by congruency effects (higher reactivation indices for past vs. future control points). For this reason, control points that are only 5 TRs (7.5 seconds) distant from the event boundaries were chosen for the 21^st^ year dataset. Since scenes in this dataset were separated by 3-4 TRs of silence, these control timepoints were the furthest possible from the event boundary while not introducing any incongruent effects into the future control point. Both control points were therefore well within a “congruent” BOLD response, thus eliminating unbalanced congruency effects. Yet this congruent response was shared with the event boundary itself. For this reason, the contrast between event boundaries and control reactivation indices, calculated for the 21^st^ year dataset, was expected to yield weaker effects as compared to effects in the Sherlock dataset.

The contrast between the reactivation index of event boundaries and control timepoints was calculated for each sphere in each participant’s brain. Single-subject and group maps were created as described previously. For consistency, a controlled reactivation analysis was additionally performed with control points that are 5 TRs away from event boundaries for the Sherlock dataset.

#### Reactivation across time

To visualize and measure the reproducibility of the observed reactivation effects across datasets (Sherlock/21^st^ year) and using the two analysis methods (within/between participants), the time-courses of reactivation were drawn. These time-courses were extracted from the previously described ROIs of brain areas that showed consistent reactivation effects across datasets: the bilateral precuneus/retrosplenial cortex, bilateral posterior hippocampus and the bilateral angular gyrus/lateral occipital cortex.

Reactivation signals from these ROIs were extracted from independent datasets and using the same analysis method: ROIs defined from the Sherlock dataset were tested on the 21^st^ year dataset, and vice versa. ROIs defined based on the within/between-participant analysis were tested using the same analysis method. This procedure allowed the conduction of 2 cross-validation tests across the independent datasets using each of the 2 analysis method.

##### Statistical analysis

For each of these ROIs, and in each participant’s data, reactivation indices were extracted for single TRs around the event boundaries, in the same manner described in the section “Reactivation analysis”. However, the current reactivation indices were uncorrected for the HRF delay, to demonstrate the true temporal correspondence between the reactivation indices and event boundaries. The time window chosen for this analysis included 3 TRs before event boundaries and 12 TRs after the event boundaries. Since fMRI autocorrelations are substantial in short time-periods (see section “Autocorrelations in within-participant analyses”), this analysis was restricted to event boundaries that originated from scenes with a minimum of 20 TRs. To provide a visual demonstration of the changes in reactivation across time, the reactivation indices were averaged across events within participant, and then averaged across participants of the same dataset.

To test the magnitude of reactivation at event boundaries across datasets and analysis methods, the following procedure was employed: the TR-specific reactivation values were divided into a baseline (the HRF-uncorrected event boundary and the TR that precedes it) and signal (TRs 1-6 after the HRF-uncorrected event boundary, to account for the HRF delay). For each participant, the averaged reactivation value of the baseline was subtracted from the averaged reactivation value of the signal. This resulted in a single reactivation index for each participant and dataset/analysis method. Note that this index is different than the original reactivation index, which measured the magnitude of reactivation at a specific TR (the HRF-corrected event boundary). Cohen’s D effect sizes were calculated by dividing each group’s mean by the groups’ standard deviation.

Group statistics were carried out using permutation tests. Single-subject indices were submitted to a one-sample t-test, and the group level t-value was defined as the statistic for the permutation test. Under the null hypothesis of no reactivation (a flat signal), the sign of single-subject reactivation indices could be flipped. Thus, for each dataset/analysis method, reactivation indices of single participants were randomly flipped, and a random group t-value was calculated. This procedure was repeated 1,000 times, resulting in 1,000 random t-values that constructed the null distribution. The position of the true test statistic in relation to the null distribution was used to determine the two-tailed p-value of the test.

To test the reproducibility of results across datasets, for each ROI separately, the 2 dataset-specific p-values resulting from the above described permutation tests were meta-analysed using Fisher’s method^65, 66^. To correct for multiple hypotheses testing across the 3 ROIs (precuneus/retrosplenial cortex, hippocampus, angular gyrus/lateral occipital cortex), the alpha level was adjusted to 0.017 based on the conservative Bonferroni correction.

#### Context-specificity of reactivation

##### whole-brain analysis

Is reactivation of remote past information context-selective? In other words, would past scenes that share a similar context with the current scene (i.e. more relevant to the event model) be reactivated more than scenes with a different context? To assess this, the full text of each experimental stimulus was analysed: descriptions of the scenes (including transcript of dialogues) in the Sherlock movie^54^, and the text of the 21^st^ year story. Under the assumption that word cooccurrences between scenes would represent context similarity, for each experimental stimulus separately, the words comprising each scene were used to construct a Bag Of Words model, as implemented in Matlab. The Bag Of Words model listed all words across each full experimental stimulus (excluding stop words, such as “the”, “a” etc., that have no specific semantic meaning), and counted the number of occurrences of each word in each scene. These scene-specific occurrence vectors were then transformed into probability vectors. Thus, for each scene, a probability vector the length of all words in the experimental narrative was created (see illustration in Figure 1d, bottom right). To measure context similarities between scenes, the Jensen-Shannon distance was calculated between each two scene-vectors, and the resulting between-scene distance matrix was converted to a similarity matrix. This matrix was meant to be correlated with the neural event-boundary X scene similarity matrix.

However, visual inspection of the Bag Of Words scene-similarity matrices of both datasets revealed a time-related bias of similarity values (sup. Figure 6a/c). Bag Of Words scene-similarities were significantly correlated with time (the serial position of scenes) in the Sherlock data (r_(901)_ = 0.69, p<0.001). This bias was related to the first 15 scenes of the movie, which were either very short (and thus had unreliable Bag Of Words representations) or less related to the overall narrative (containing non-central characters/events, which resulted in unique Bag Of Word representations). The first 15 scenes were indeed less similar to all other scenes, compared to the between-scene similarities of the remaining scenes (t_(901)_ = -25.9, p<0.001, sup. Figure 6a). Thus, correlating the Bag Of Words scene-similarity matrix with the neural similarity matrix would capture time-related effects, rather than the more subtle modulation of reactivation by semantic context. A similar time-related bias was apparent across the congruent entries of the 21^st^ year Bag Of Words scene-similarity matrix (r_(763)_ = - 0.08, p=0.03). This bias was related to the merging of the two separate narratives in the last 15 scenes of the story. By definition, the congruent entries of each separate narrative would be more similar to each other (as they contain very similar words) compared to the similarities between the entries of the joint narrative and each of the separate narratives (since the joint narrative contains a mixture of words from both narratives, and would therefore be less similar to each separate narrative). The between-scene similarities of the last 15 scenes (which are by definition congruent with all other scenes) were indeed lower compared to the congruent entries of the between-scene similarities of the first 30 scenes (t_(553)_ = -10.22, p<0.001, sup. Figure 6c). To prevent these time-related biases from masking true experimental effects, the first 15 scenes of the Sherlock dataset and the last 15 scenes of the 21^st^ year dataset were excluded from the context-similarity matrices. This procedure successfully eliminated the time biases (r_(433)_ = 0.05/0.02, p = 0.3/0.7, for Sherlock/21^st^ year, sup. Figure 6b/d, respectively). For transparency, a whole-brain Bag Of Words analysis was also conducted using the full, time-biased Bag Of Words context-similarity matrices.

Thus, for the Sherlock dataset, the following analysis was performed: for each participant and sphere, a neural event-boundary X scene similarity matrix was computed using either a within- or between-participants analysis, as previously described. A context scene-similarity matrix was also calculated. Due to the variable scene-length in the Sherlock movie (Table 1), which could affect the context similarity values of short scenes (with very sparse vectors), a third matrix indicating the minimal number of words in each scene-pair was also created. Next, the past part of the neural similarity matrix was correlated with the past part of the context similarity matrix, and the future part of the neural similarity matrix was correlated with the future part of the context similarity matrix (which is symmetric to the past part of the same matrix). In both computations, the contribution of the minimal-word matrix was partialled out of the correlation. Finally, the difference between the Fisher-transformed past and future correlation coefficients was computed, cancelling out any effect of scene-similarities, which is symmetric in time. This difference value was assigned to the center of the sphere. Single-subject and group maps were created as described for the reactivation analysis. However, given the clear directionality of the experimental hypothesis (similarity in context will induce more reactivation), one-tailed p-values were computed.

The same analysis could not be performed for the 21^st^ year dataset, due to its experimental design. Due to the alternating narratives across the first 30 scenes, a checkerboard pattern of similarity was captured by the Bag Of Words model (sup. Figure 6c/d). Similarly to the previously described time-bias, correlating the context scene-similarity matrix with the neural similarity matrix would capture the robust congruency effect, rather than the more subtle modulation of reactivation by sematic context in the congruent scenes. For this reason, rather than using the main effect of time as the basis for the neural reactivation index, a time X congruency interaction was used. Specifically, for each participant and sphere, event-boundary X scene similarity matrices were calculated, as previously described. Next, separately for the past and future parts of each neural similarity matrix, the congruent entries were correlated with the corresponding entries in the context similarity matrix, and the incongruent entries were correlated with corresponding entries in the context similarity matrix. Finally, the difference between the Fisher-transformed congruent and incongruent correlations of the past part of the matrix were contrasted with the difference between the Fisher-transformed congruent and incongruent correlations of the future part of the matrix. This index was assigned to the center of each sphere. Single-subject and one-tailed group maps were created as described for the reactivation analysis.

##### ROI analysis

To test whether the same regions that showed a reactivation effect also showed modulation of this reactivation by semantic context, an ROI analysis was performed. This analysis was confined to the precuneus/retrosplenial cortex, since this was the only region that showed a reproducible context modulation effect across datasets and analysis methods in the whole-brain analysis. The same precuneus/retrosplenial cortex ROIs that were defined from the reactivation analysis group maps were used. Since these ROIs were based on the reactivation analysis, they were independent with regards to context modulation analysis. Thus, the ROIs defined from the reactivation analysis for each dataset (Sherlock/21^st^ year) and analysis method (within/between-participants) were used to test the modulation of reactivation by semantic context for the same dataset and analysis method. Specifically, for each participant, the relevant (dataset- and analysis-specific) ROI was used to extract event-boundaries and scene representations. Neural similarity matrices were constructed, and correlated with a context similarity matrix, as previously described. This resulted in 4 sets of values derived from single-participants (Sherlock/21^st^ year participants in the within/between-participants analysis).

Group statistics were carried out using permutation tests. Each set of single-subject values was submitted to a one-sample t-test, and the group level t-value was defined as the statistic for the permutation test. Under the null hypothesis of no contextual modulation of reactivation, the sign of single-subject values could be flipped. Thus, for each dataset/analysis method, values of single participants were randomly flipped, and a random group t-value was calculated. This procedure was repeated 1,000 times, resulting in 1,000 random t-values that constructed the null distribution. The position of the true test statistic in relation to the null distribution was used to determine the one-tailed p-value of the test.

To test the reproducibility of the contextual modulation of reactivation in the precuneus/retrosplenial cortex, the 2 dataset-specific p-values resulting from the above described permutation tests were meta-analysed using Fisher’s method. This was done separately for each of the analysis methods (within/between participants).

#### Mega analyses

A pooled whole-brain analysis across datasets was conducted using FSL’s randomise. Single-subject maps of all participants were mega-analysed using a second-level GLM design, where each dataset was modelled with a dummy regressor. Resulting mega-analysis maps were thresholded and corrected for multiple comparisons, as described for the within-dataset group analyses.

A mega-analysis of the context-specificity of reactivation in the precuneus/retrosplenial cortex was also conducted using a permutation test. The only difference to the permutation test previously described for the within-dataset analysis was the inclusion of dummy regressors for each dataset.

#### Autocorrelations in within-participant analyses

Calculating correlations between representations of single TRs (event boundaries) and other timepoints of the same fMRI signals is problematic, since these representations are embedded in 1/f noise (slow trends in the time domain, which contribute to the lower frequencies of the power spectrum)^67^. This noise is typically removed by applying a linear high-pass filter, which removes from each TR the commonalities with its weighted temporal neighbourhood. By doing so, the correlations between each TR and its environment are removed, thus “flattening” the linear trend induced by the 1/f noise (sup. Figure 8a). However, this procedure changes the autocorrelation structure of fMRI data: prior to filtering, each TR has positive correlations with neighbouring TRs that slowly decline in time, but high-pass filtering induces a more complex structure of alternating positive and negative correlations that decline in time (sup. Fig. 8b). Thus, correlations between event boundaries and scenes will have a temporal structure, which is symmetric in time, because event boundaries are symmetric with regards to past and future scenes. However, since the scenes that immediately precede event boundaries are removed from the similarity matrices, past scenes are further away from event boundaries compared to future scenes. This entails that the temporal structure introduced by autocorrelations will be asymmetric with respect to past and future scenes, thus biasing the reactivation index.

Moreover, this temporal bias will also interact with any temporal asymmetries introduced by the structure of the experimental stimulus. Datasets based on experimental stimuli with short scenes (e.g. Sherlock dataset) will be more heavily impacted by autocorrelation biases compared to datasets based on experimental stimuli with longer scenes (e.g. 21^st^ year dataset). The relative position of short scenes will also determine the magnitude of bias (series of adjacent short scenes will be heavily impacted). For example, the beginning of the movie Sherlock has a cluster of short scenes, and therefore, on average, event boundaries will be closer in time to past scenes than to future scenes. In addition, inter-scene intervals, such as those in the 21^st^ year story, entail that event boundaries are closer to their past (containing) scenes, compared to the future (non-continuous) scenes. Finally, datasets based on experimental stimuli with a time-alternating structure (e.g. 21^st^ year dataset) will have a different bias compared to datasets devoid of such a structure (e.g. Sherlock dataset), as will be discussed below. Therefore, to determine what would be the most suitable unbiased statistic for each dataset (Sherlock/21^st^ year), simulations based on the unique characteristics of each dataset and its related experimental stimulus were employed.

Note again that these autocorrelation biases do not exist in the between-participant analysis, where the representations of event boundaries and scenes are extracted from different brains (independent signals). The fact that the between-participant effects cleanly replicate the within-participant effects should increase confidence that autocorrelations have been suitably accounted for using the approach described in the following sections.

#### Reactivation index

To control for the autocorrelation bias, adjacent scenes on both sides of event boundaries, which are heavily impacted by the bias, must be remove from our analyses. To assess the number of scenes that must be removed, 10 “single subject” simulated datasets of 120 voxels with a 1/f power spectrum were created and high-pass filtered. These datasets were created based on either the characteristics of the Sherlock experiment (TR=1.5, scan length=1,976 TRs, high-pass filter of 140 sec) or the 21^st^ year experiment (TR=1.5, scan length=2,242 TRs, high-pass filter of 120 sec). In each simulated dataset, representations of event boundaries and scenes were extracted based on the specific timings of scenes in the experimental stimuli. Representations of event boundaries and scenes were then correlated within each simulated dataset, yielding 10 random similarity matrices. These 10 matrices were used to calculate 10 random reactivation indices, which were then averaged into a group index. The 10 similarity matrices were later averaged across the simulated datasets to create a group similarity matrix. This procedure was repeated 10,000 times, resulting in 10,000 group reactivation indices and 10,000 group similarity matrices. The group similarity matrices were further averaged for display purposes.

As evident in the upper left panels of sup. Figure 8c/d, both the averaged Sherlock and 21^st^ year similarity matrices showed artefactual correlations around their main diagonals (the missing rows/columns in the Sherlock matrix correspond to excluded scenes and event boundaries, as described above). The lower left panels of sup. Figure 8c/d demonstrate that these artefactual correlations were reflected in positively biased distributions of the reactivation indices. However, removing 10 diagonals from the random Sherlock matrices and 6 diagonals from the random 21^st^ year matrices removed the artefactual correlations (upper right panels of sup. Figure 8c/d) and centred the reactivation index distributions on 0 (lower right panels of sup. Figure 8c/d), thus removing autocorrelation-related biases from our analysis.

Note that removing diagonals from the similarity matrix means discarding of scenes that precede or follow each event boundary (5/3 scenes on each side of event boundaries in the Sherlock/21^st^ year datasets, respectively). An equal number of diagonals from the past and future parts of the similarity matrix must be discarded, since high-pass filtering has a symmetric effect on timepoints that precede of follow each event boundary. For this same reason, the main diagonal must belong to the past part of the similarity matrix, since it contains the scenes immediately preceding each event boundary. Removal of n diagonals from the similarity matrix therefore entails the removal of n/2 diagonals from the upper triangular part of the matrix (future part), and the removal of the main diagonal and additional n/2-1 diagonals from the lower part of the matrix (past part). The reactivation index would therefore take the following form:

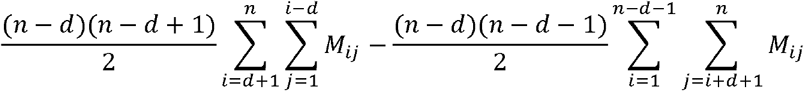

Where M is an n X n matrix, i and j are row and column indices, respectively, and d is the number of diagonals removed from the upper triangular part of the matrix.

Though our analyses create a slightly unbalanced partition of the similarity matrix (more past than future entries), the simulations presented in the lower right panels of sup. Figure 8c/d demonstrate that this imbalance does not bias the reactivation index distribution.

To assess whether stable scene patterns may also introduce some bias into the analysis, the simulations described above were repeated while adding in random scene representations drawn from a uniform distribution. These added representations were given weights of 0.1, 0.5, 1 and 1.5 with regards to the averaged scene representations derived from the random 1/f signals. As demonstrated in sup. Figure 9 (for weights 0.5 and 1), the higher the weight, the more the added patterns masked the autocorrelation bias, as scene representations no longer reflected solely the 1/f signals they arose from. Yet the removal of diagonals was still needed to center the reactivation index distribution on zero. In other words, these uncorrelated scene patterns simply added unexplained variance to the analysis, suggesting that it is clearly not mere scene representations that cause the observed bias.

To investigate whether autocorrelations between scene representations could cause the bias, simulations were repeated while adding in weighted scene patterns drawn from a 1/f distribution. As demonstrated in sup. Figure 10, adding in autocorrelated scene representations did not bias the analysis, but only masked the time bias, as scene patterns no longer solely reflect the originating 1/f signals (and again, removal of diagonals was needed to control for the signal autocorrelation bias). This is because scene autocorrelations are a function of scene order and not of time. This means they are perfectly symmetric with regards to event boundaries and will get subtracted out by our measure.

To assess whether a congruency structure, as in the 21^st^ year dataset, could underlie the bias, random template representations were drawn for narrative A and for narrative B of the story. These representations were added to the simulated scene representations in an interleaved manner, and were added gaussian noise to create correlations between congruent scene representations. As demonstrated in sup. Figure 11, adding in these weighted scene patterns again masked the observed bias, overruling the possibility that mere congruency structure caused the autocorrelation bias.

However, the congruency structure of the 21^st^ year dataset does interact with the time-related autocorrelation bias, which prevented the use of a Time (past/future) X Congruency (same/different narrative) interaction in the calculation of the reactivation index. As previously described, and schematically illustrated in sup. Figure 12a, the first 30 scenes of the 21^st^ year datasets consisted of alternating narratives, that later converge. The similarity matrices calculated for this dataset would therefore contain alternating entries of congruent (same narrative) or incongruent (different narrative) correlations between event boundaries and scenes. These entries are arranged by diagonals, which create an asymmetric congruency structure with regards to event boundaries. For example, the main diagonal contains correlations between event boundaries and their immediately preceding scenes, which are congruent. The superdiagonal contains correlations between event boundaries and their immediately following scenes, which are incongruent. However, autocorrelations have a symmetric effect in time. Event boundaries are more correlated to closer points in time, and are hence more correlated to congruent scenes in the past and to incongruent scenes in the future, which will bias any congruency-based analysis, such as a Time X Congruency interaction. This bias cannot be fixed by removing diagonals from the analysis, since an equal number of diagonals must be removed from the past/future parts of the matrix, which will maintain an asymmetric congruency structure in time. Indeed, when repeating the simulations of our analysis with random 1/f data, as described above, but this time defining the reactivation index as a Time X Congruency interaction, the removal of the diagonals failed to remove the autocorrelation artefact from the reactivation index distribution (the distribution is positively biased, sup. Figure 12b). For this reason, a reactivation index that captures the main effect of time (past-future) was chosen for both the Sherlock and 21^st^ year datasets. For consistency, the same reactivation index was also used in the main between-participant analyses.

##### Control time-points

As explained in the section “Time-specificity of reactivation”, the choice of an appropriate time-point to serve as a control is non-trivial, due to the influence of autocorrelations. Event boundaries are by definition positioned between two adjacent scenes. Due to this position, the effect of high-pass filtering on the event boundary is symmetric with respect to time, since the local temporal environment of the event boundary contains equal time-points from past and future scenes (this is true for the Sherlock dataset, where no time-gaps exist between scenes. This is marginally true for the 21^st^ year dataset, where inter-scene intervals were inserted, making event boundaries slightly closer to past scenes than to future scenes). However, a control time-point that precedes an event boundary will be much closer to the past period (as defined by the boundary) and a control time-point that follows an event boundary will be much closer to the future period (sup. Figure 13a). In both cases, high-pass filtering will induce unbalanced temporal autocorrelations between the control point and past/future scenes, and will therefore introduce a temporal bias into our analyses.

To illustrate this point, simulations of random high-pass filtered 1/f data were repeated for the two experiments, as described above, but this time, past (10 timepoints before event boundaries) and future (10 timepoints after event boundaries) control timepoints were added to the analysis. For each of these timepoints separately (event boundaries, past/future controls), representations of the relevant timepoints were extracted. These representations were then correlated with scene representations to construct similarity matrices, diagonals were discarded as described above, and reactivation indices were computed. As depicted in sup. Figure 13b, and as previously demonstrated, the distribution of the reactivation index at event boundaries was centred on zero in the datasets of both experiments. However, the distributions of reactivation indices of both the past and future control points were not centered on zero, indicating a high-pass-filtering artefact that interacts with time (sup. Figure 13c/d). The results of these simulations indicate that contrasting the reactivation indices of the event boundaries with those of either past or future control timepoints would introduce biases into our analyses. However, since high-pass filtering is symmetric in time, using both control time points, which are equally distant from the event boundary, would cancel out the temporal bias and create a balanced control. Simulation analyses indeed revealed that contrasting the reactivation index of event boundaries with that of the balanced control yielded distributions centered on zero (sup. Figure 13e), indicating an unbiased analysis.

##### Context-modulation of reactivation analysis in the 21^st^ year dataset

As previously demonstrated, using a time X congruency interaction-based reactivation index for the 21^st^ year dataset inherently induced biases into the analysis. To evaluate whether the context-modulation analysis would be similarly affected by this bias, simulations were performed. In these simulations, the interaction of correlations between the random neural similarity matrices and the context similarity matrices was computed. This procedure was repeated 1,000 times, constructing a random distribution of interaction values, which showed a negative bias (sup. Figure 12c). Since this bias was in the opposite direction to the experimental hypothesis, it would attenuate rather than enhance any experimental effect. Thus, weaker results were expected to be found for this analysis in the 21^st^ year dataset compared to the Sherlock dataset.

## Code availability

Matlab code will be uploaded to GitHub upon publication.

## References

1. Zacks, J.M., Speer, N.K., Swallow, K.M., Braver, T.S. & Reynolds, J.R. Event perception: a mind-brain perspective. Psychological bulletin 133, 273 (2007).

2. Radvansky, G.A. & Zacks, J.M. Event perception. Wiley Interdisciplinary Reviews: Cognitive Science 2, 608–620 (2011).

3. Zwaan, R.A. & Radvansky, G.A. Situation models in language comprehension and memory. Psychological bulletin 123, 162 (1998).

4. Bird, C.M. How do we remember events? Current Opinion in Behavioral Sciences 32, 120–125 (2020).

5. Stawarczyk, D., Bezdek, M.A. & Zacks, J.M. Event representations and predictive processing: The role of the midline default network core. Topics in Cognitive Science 13, 164–186 (2021).

6. Tolman, E.C. Cognitive maps in rats and men. Psychol Rev 55, 189–208 (1948).

7. Eichenbaum, H. A cortical–hippocampal system for declarative memory. Nature reviews neuroscience 1, 41–50 (2000).

8. Holland, P.C. Acquisition of representation-mediated conditioned food aversions. Learning and Motivation 12, 1–18 (1981).

9. Chen, J., et al. Accessing real-life episodic information from minutes versus hours earlier modulates hippocampal and high-order cortical dynamics. Cerebral cortex 26, 3428–3441 (2016).

10. Yeshurun, Y., Nguyen, M. & Hasson, U. The default mode network: where the idiosyncratic self meets the shared social world. Nature Reviews Neuroscience 22, 181–192 (2021).

11. Keidel, J.L., Oedekoven, C.S., Tut, A.C. & Bird, C.M. Multiscale integration of contextual information during a naturalistic task. Cerebral Cortex 28, 3531–3539 (2018).

12. Cohn-Sheehy, B.I., et al. The hippocampus constructs narrative memories across distant events. Current Biology 31, 4935-4945. e4937 (2021).

13. Foster, D.J. Replay comes of age. Annual review of neuroscience 40, 581–602 (2017).

14. Findlay, G., Tononi, G. & Cirelli, C. The evolving view of replay and its functions in wake and sleep. Sleep Advances 1, zpab002 (2020).

15. Foster, D.J. & Wilson, M.A. Reverse replay of behavioural sequences in hippocampal place cells during the awake state. Nature 440, 680–683 (2006).

16. Singer, A.C. & Frank, L.M. Rewarded outcomes enhance reactivation of experience in the hippocampus. Neuron 64, 910–921 (2009).

17. Silva, M., Baldassano, C. & Fuentemilla, L. Rapid memory reactivation at movie event boundaries promotes episodic encoding. Journal of Neuroscience 39, 8538–8548 (2019).

18. Sols, I., DuBrow, S., Davachi, L. & Fuentemilla, L. Event boundaries trigger rapid memory reinstatement of the prior events to promote their representation in long-term memory. Current Biology 27, 3499-3504. e3494 (2017).

19. Ben-Yakov, A., Eshel, N. & Dudai, Y. Hippocampal immediate poststimulus activity in the encoding of consecutive naturalistic episodes. Journal of Experimental Psychology: General 142, 1255 (2013).

20. Ben-Yakov, A. & Henson, R.N. The Hippocampal Film Editor: Sensitivity and Specificity to Event Boundaries in Continuous Experience. J Neurosci 38, 10057–10068 (2018).

21. Raichle, M.E., et al. A default mode of brain function. Proc Natl Acad Sci U S A 98, 676–682 (2001).

22. Speer, N.K., Zacks, J.M. & Reynolds, J.R. Human brain activity time-locked to narrative event boundaries. Psychological Science 18, 449–455 (2007).

23. Zacks, J.M., Speer, N.K., Swallow, K.M. & Maley, C.J. The brain’s cutting-room floor: Segmentation of narrative cinema. Frontiers in human neuroscience 4, 168 (2010).

24. Whitney, C., et al. Neural correlates of narrative shifts during auditory story comprehension. Neuroimage 47, 360–366 (2009).

25. Abadchi, J.K., et al. Spatiotemporal patterns of neocortical activity around hippocampal sharp-wave ripples. Elife 9, e51972 (2020).

26. Higgins, C., et al. Replay bursts in humans coincide with activation of the default mode and parietal alpha networks. Neuron 109, 882-893. e887 (2021).

27. Norman, Y., Raccah, O., Liu, S., Parvizi, J. & Malach, R. Hippocampal ripples and their coordinated dialogue with the default mode network during recent and remote recollection. Neuron 109, 2767-2780. e2765 (2021).

28. Kaplan, R., et al. Hippocampal sharp-wave ripples influence selective activation of the default mode network. Current Biology 26, 686–691 (2016).

29. Pedrosa, R., et al. Hippocampal gamma and sharp wave/ripples mediate bidirectional interactions with cortical networks during sleep. bioRxiv (2022).

30. Tambini, A. & Davachi, L. Awake reactivation of prior experiences consolidates memories and biases cognition. Trends in cognitive sciences 23, 876–890 (2019).

31. Gupta, A.S., van der Meer, M.A., Touretzky, D.S. & Redish, A.D. Hippocampal replay is not a simple function of experience. Neuron 65, 695–705 (2010).

32. Pfeiffer, B.E. & Foster, D.J. Hippocampal place-cell sequences depict future paths to remembered goals. Nature 497, 74–79 (2013).

33. Karlsson, M.P. & Frank, L.M. Awake replay of remote experiences in the hippocampus. Nat Neurosci 12, 913–918 (2009).

34. Liu, Y., Dolan, R.J., Kurth-Nelson, Z. & Behrens, T.E. Human replay spontaneously reorganizes experience. Cell 178, 640-652. e614 (2019).

35. Schwartenbeck, P., et al. Generative replay for compositional visual understanding in the prefrontal-hippocampal circuit. bioRxiv (2021).

36. Kriegeskorte, N., Mur, M. & Bandettini, P. Representational similarity analysis–connecting the branches of systems neuroscience. Frontiers in systems neuroscience 2 (2008).

37. Chen, J. Sherlock Movie Watching Dataset. (2016).

38. Nastase, S., et al. The “Narratives” fMRI dataset for evaluating models of naturalistic language comprehension. Scientific Data, 8 (1), 250–250. (2021).

39. Chang, C.H., Lazaridi, C., Yeshurun, Y., Norman, K.A. & Hasson, U. Relating the past with the present: Information integration and segregation during ongoing narrative processing. Journal of Cognitive Neuroscience 33, 1106–1128 (2021).

40. Chen, J., et al. Shared memories reveal shared structure in neural activity across individuals. Nat Neurosci 20, 115–125 (2017).

41. Zadbood, A., Chen, J., Leong, Y.C., Norman, K.A. & Hasson, U. How we transmit memories to other brains: constructing shared neural representations via communication. Cerebral cortex 27, 4988–5000 (2017).

42. Chien, H.-Y.S. & Honey, C.J. Constructing and forgetting temporal context in the human cerebral cortex. Neuron 106, 675-686. e611 (2020).

43. Pfeiffer, B.E. The content of hippocampal “replay”. Hippocampus 30, 6–18 (2020).

44. Buckner, R.L., Andrews-Hanna, J.R. & Schacter, D.L. The brain’s default network: anatomy, function, and relevance to disease. Ann N Y Acad Sci 1124, 1–38 (2008).

45. Buckner, R.L. & Carroll, D.C. Self-projection and the brain. Trends in cognitive sciences 11, 49–57 (2007).

46. Yeshurun, Y., et al. Same story, different story: the neural representation of interpretive frameworks. Psychological science 28, 307–319 (2017).

47. Baldassano, C., et al. Discovering event structure in continuous narrative perception and memory. Neuron 95, 709-721. e705 (2017).

48. Stephens, G.J., Honey, C.J. & Hasson, U. A place for time: the spatiotemporal structure of neural dynamics during natural audition. Journal of neurophysiology 110, 2019–2026 (2013).

49. Shohamy, D. & Wagner, A.D. Integrating memories in the human brain: hippocampal-midbrain encoding of overlapping events. Neuron 60, 378–389 (2008).

50. Schlichting, M.L., Zeithamova, D. & Preston, A.R. CA1 subfield contributions to memory integration and inference. Hippocampus 24, 1248–1260 (2014).

51. Collin, S.H., Milivojevic, B. & Doeller, C.F. Memory hierarchies map onto the hippocampal long axis in humans. Nature neuroscience 18, 1562–1564 (2015).

52. Eichenbaum, H. The hippocampus and mechanisms of declarative memory. Behavioural brain research 103, 123–133 (1999).

53. Miller, A.M., Vedder, L.C., Law, L.M. & Smith, D.M. Cues, context, and long-term memory: the role of the retrosplenial cortex in spatial cognition. Frontiers in human neuroscience 8, 586 (2014).

54. Lee, H. & Chen, J. Narratives as Networks: Predicting Memory from the Structure of Naturalistic Events. bioRxiv (2021).

55. Singer, A.C., Carr, M.F., Karlsson, M.P. & Frank, L.M. Hippocampal SWR activity predicts correct decisions during the initial learning of an alternation task. Neuron 77, 1163–1173 (2013).

56. Jai, Y.Y. & Frank, L.M. Hippocampal–cortical interaction in decision making. Neurobiology of learning and memory 117, 34–41 (2015).

57. Gillespie, A.K., et al. Hippocampal replay reflects specific past experiences rather than a plan for subsequent choice. bioRxiv (2021).

58. Jenkinson, M., Bannister, P., Brady, M. & Smith, S. Improved optimization for the robust and accurate linear registration and motion correction of brain images. Neuroimage 17, 825–841 (2002).

59. Jenkinson, M. & Smith, S. A global optimisation method for robust affine registration of brain images. Med Image Anal 5, 143–156 (2001).

60. Greve, D.N. & Fischl, B. Accurate and robust brain image alignment using boundary-based registration. Neuroimage 48, 63–72 (2009).

61. Hahamy, A., et al. Representation of Multiple Body Parts in the Missing-Hand Territory of Congenital One-Handers. Curr Biol 27, 1350–1355 (2017).

62. Oedekoven, C.S., Keidel, J.L., Berens, S.C. & Bird, C.M. Reinstatement of memory representations for lifelike events over the course of a week. Scientific Reports 7, 1–12 (2017).

63. Bird, C.M., Keidel, J.L., Ing, L.P., Horner, A.J. & Burgess, N. Consolidation of complex events via reinstatement in posterior cingulate cortex. Journal of Neuroscience 35, 14426–14434 (2015).

64. Hasson, U., Nir, Y., Levy, I., Fuhrmann, G. & Malach, R. Intersubject synchronization of cortical activity during natural vision. Science 303, 1634–1640 (2004).

65. Fisher, R.A. Statistical methods for research workers (Genesis Publishing, 1925).

66. Fisher, R. Questions and answers# 14. The American Statistician 2, 30–31 (1948).

67. Zarahn, E., Aguirre, G. & DEsposito, M. Empirical analyses of BOLD fMRI statistics. 1. Spacially unsmoothed data collected under null-hypothesis conditions (vol 5, pg 179, 1997). 71-72 (ACADEMIC PRESS INC JNL-COMP SUBSCRIPTIONS 525 B ST, STE 1900, SAN DIEGO, CA …, 1997).

