## Supplementary for "The human brain reactivates context-specific past information at event boundaries of naturalistic experiences"

| ROI definition | Test | PCUN | Hippocampus | Ang/LOC |
|---|---|---|---|---|
| Sherlock<br>within-participant | 21st year<br>within-participant | M=0.018<br>SEM=0.004<br>D=0.84<br>p<0.001 | M=0.006<br>SEM=0.004<br>D=0.34<br>p=0.044 | M=0.007<br>SEM=0.003<br>D=0.42<br>p=0.02 |
| 21st year<br>within-participant | Sherlock<br>within-participant | M=0.039<br>SEM=0.01<br>D=0.96<br>p<0.001 | M=0.01<br>SEM=0.01<br>D=0.28<br>p=0.1 | M=0.048<br>SEM=0.01<br>D=1.2<br>p<0.001 |
| Sherlock<br>between-participants | 21st year<br>between-<br>participants | M=0.018<br>SEM=0.004<br>D=0.83<br>p<0.001<br><br>(M=0.02<br>SEM=0.006<br>D=0.62<br>p<0.001) | M=0.01<br>SEM=0.002<br>D=0.83<br>p=0.001<br><br>(M=0.01<br>SEM=0.005<br>D=0.42<br>p=0.001) | M=0.012<br>SEM=0.002<br>D=1.08<br>p<0.001<br><br>(M=0.01<br>SEM=0.004<br>D=0.79<br>p<0.001) |
| 21st year<br>between-participants | Sherlock<br>between-<br>participants | M=0.032<br>SEM=0.008<br>D=0.99<br>p<0.001<br><br>(M=0.04<br>SEM=0.007<br>D=1.56<br>p<0.001) | M=0.04<br>SEM=0.014<br>D=0.68<br>p=0.008<br><br>(M=0.07<br>SEM=0.02<br>D=1.08<br>p=0.008) | M=0.054<br>SEM=0.01<br>D=1.05<br>p<0.001<br><br>(M=0.09<br>SEM=0.01<br>D=1.86<br>p=0.003) |

**Supplementary Table 1. Statistical parameters for the independent ROI analyses of the reactivation index, corresponding to Figure 4 and Supplementary Figures 5.**
Each row depicts the results of ROI analyses based on the independent datasets and the two analysis methods (within/between-participant reactivation). The first column specifies the dataset and analysis method used to define each ROI. The second column specifies the independent dataset in which these ROIs were tested, using the same analysis method. For the between-participants analysis, main statistics are derived from the boundary-based approach, while statistics in parentheses are derived from the scene-based approach. All p-values are two-tailed. PCUN, precuneus/retrosplenial cortex; Ang/LOC, Angular gyrus/Lateral Occipital Cortex; M, mean; D, Cohen's D.

| ANALYSIS METHOD | SHERLOCK | 21$^{ST}$ YEAR |
|---|---|---|
| WITHIN-PARTICIPANT | M=0.11<br>SEM=0.19<br>D=1.45 | M=0.054<br>SEM=0.029<br>D=0.38 |
| BOUNDARY-BASED BETWEEN-PARTICIPANT | M=0.099<br>SEM=0.016<br>D=1.48 | M=0.046<br>SEM=0.031<br>D=0.3 |
| SCENE-BASED BETWEEN-PARTICIPANT | M=0.11<br>SEM=0.01<br>D=2.56 | M=0.065<br>SEM=0.027<br>D=0.5 |

**Supplementary Table 2. Means (M), SEMs and Cohen's effect sizes (D) for the Bag Of Words independent ROI analyses, corresponding to Figure 5 and Supplementary Figures 7.**

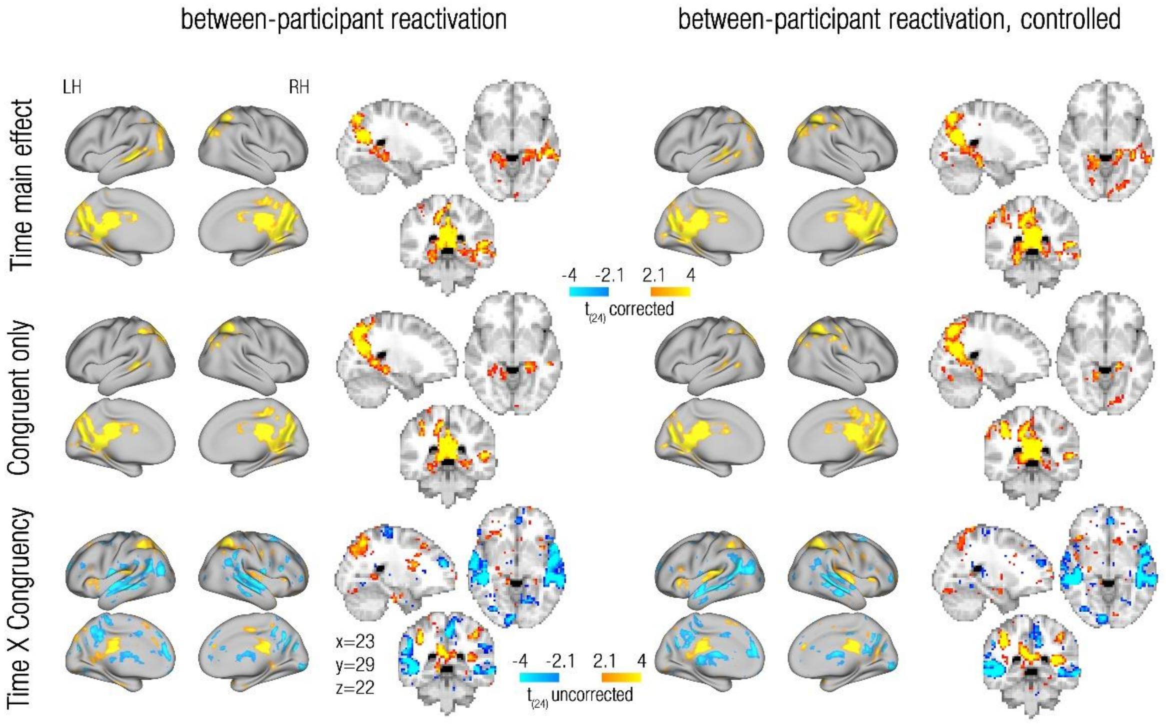


**Supplementary Figure 1. Between-participant reactivation, congruency-related analyses in the 21st year dataset**

**(a)** Between-participants reactivation (uncontrolled on the left, controlled on the right), where reactivation indices were calculated as the time main effect (difference between the mean past entries and mean future entries of the neural correlation matrix), as presented in Figure 3. This calculation included both the congruent and incongruent entries of the neural correlation matrix. Maps were cluster-corrected for multiple comparisons across the entire brain ($p<0.005$) **(b)** Between-participants reactivation, where reactivation indices were calculated as the time main effect of the congruent entries of the past and future parts of the neural correlation matrix. Maps were cluster-corrected for multiple comparisons across the entire brain ($p<0.005$). Note the resemblance to the maps presented in (a). **(c)** Between-participants reactivation, where reactivation indices were calculated as the Time (past/future) X Congruency (same/different narrative) interaction. These interaction maps did not survive correction for multiple comparisons. Maps are presented on both flat cortical surface and on 3D slices (MNI coordinates are depicted at the bottom). LH, left hemisphere; RH, right hemisphere.

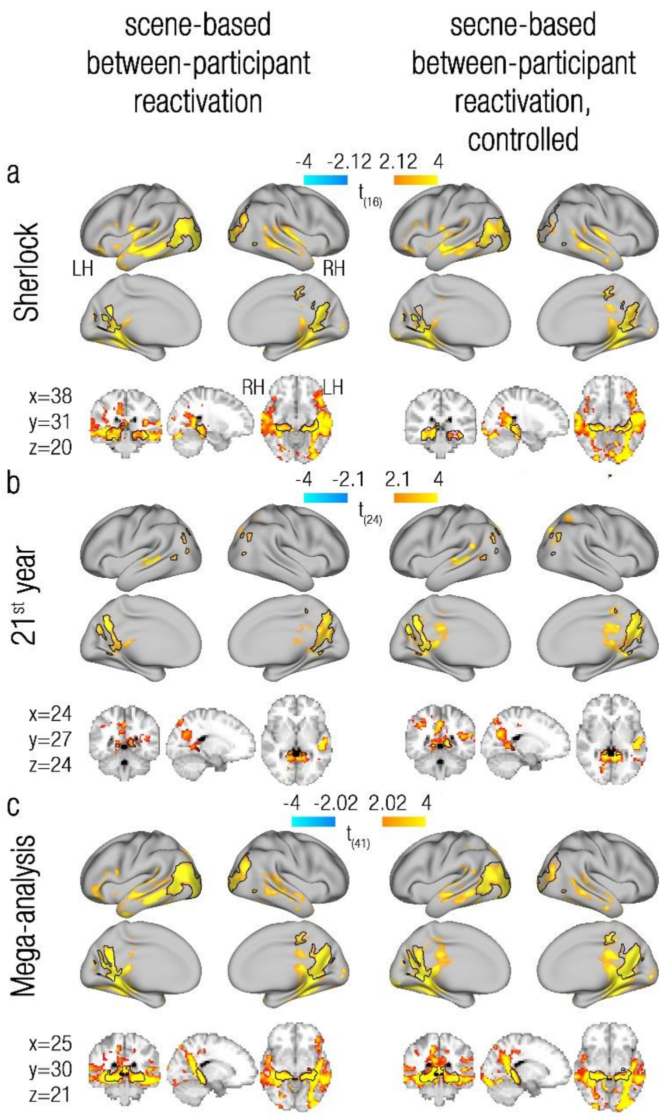


**Supplementary Figure 2. Reactivation of remote past events at event boundaries: scene-based between-participant analysis**

A scene-based between-participants reactivation analysis, where reactivation indices were derived from the event boundary representations of one participant and scene representations from all other participants of the same dataset (1st column); and a controlled scene-based between-participant reactivation analysis, where the between-participant reactivation indices at event boundaries were contrasted with those computed for control timepoints (2nd column). Rows show within-dataset group maps of the Sherlock dataset **(a)**, the 21st year dataset **(b)**, and a mega-analysis pooling together the two datasets **(c)**. All maps are projected onto a flat cortical surface, and subcortical results are also presented using 3D brain slices (dataset-specific coordinates are given at the bottom left). Similar to the within-participant analysis (Figure 2) and to the boundary-based between-participant analysis (Figure 3), reactivation of remote past events (as reflected in positive t-values) was consistently found in the two datasets in the bilateral precuneus/retrosplenial cortex (PCUN), posterior hippocampus (Hip) and the Angular gyrus/Lateral Occipital Cortex (Ang/LOC). These ROIs were defined based on the uncontrolled analysis of each dataset, as marked in black contours, and superimposed on the corresponding controlled reactivation map. LH, left hemisphere; RH, right hemisphere. All maps were cluster-corrected for multiple comparisons across the entire brain ($p<0.005$).

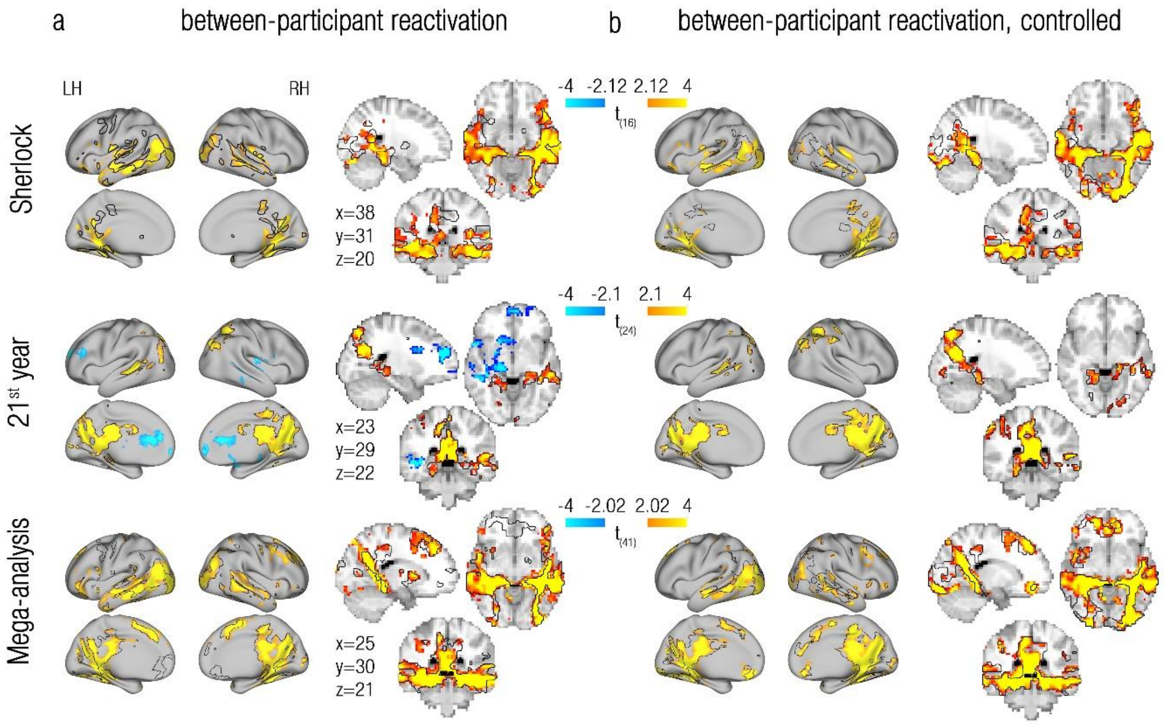


**Supplementary Figure 3. Between-participant reactivation, controlled for potential flushed scene representations**

**(a)** Between-participants reactivation, where reactivation indices were derived from the scene representations of one participant and the event boundary representations from all other participants of the same dataset. **(b)** Controlled between-participant reactivation, where the between-participant reactivation indices at event boundaries were contrasted with those computed for control timepoints. Rows depict within-dataset group maps of the Sherlock dataset (1$^{st}$ row), the 21$^{st}$ year dataset (2$^{nd}$ row) and a mega-analysis pooling together the two datasets (3$^{rd}$ row). All maps were produced using a control analysis that discards the superdiagonal. The contours of the original maps, presented in Figure 3, are superimposed over these control maps. Maps are presented on both flat cortical surface and on 3D slices (dataset-specific MNI coordinates are depicted). LH, left hemisphere; RH, right hemisphere. All maps were cluster-corrected for multiple comparisons across the entire brain (p<0.005).

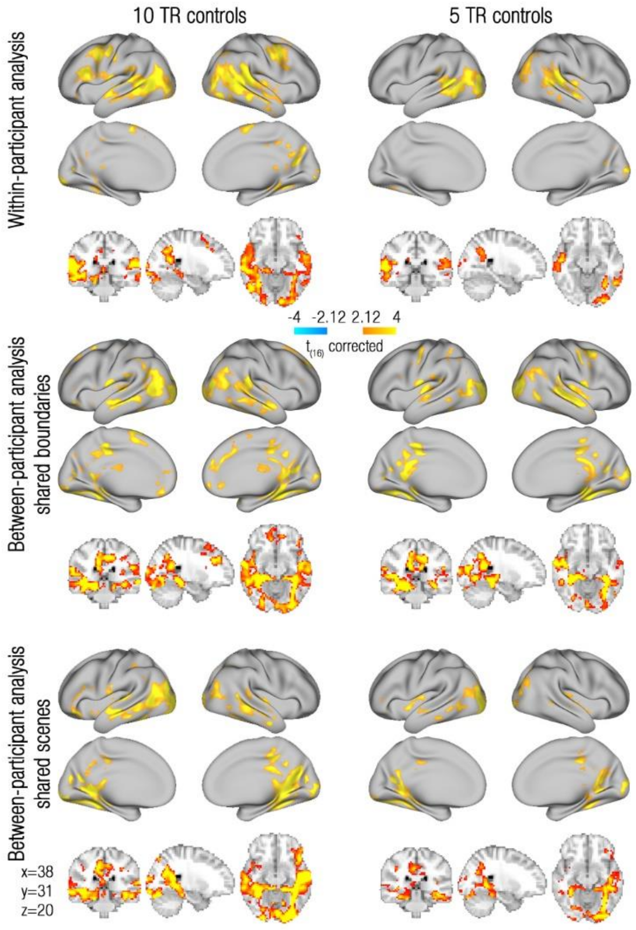


**Supplementary Figure 4. Controlled reactivation analysis in the Sherlock dataset using control points that are either 5 or 10 TRs away from event boundaries.**

Rows depict analysis methods: within-participant analysis (1st row), boundary-based between-participant analysis (2nd row), and scene-based between-participant analysis (3rd row). Maps based on a 10 TR control (left column) are the same as in Figure 2, Figure 3 and sup. Figure 2, and are presented here as reference for the maps that are based on a 5 TR control (right column). All maps are projected onto a flat cortical surface, and subcortical results are also presented using 3D brain slices (coordinates are given on the bottom left). All maps were cluster-corrected for multiple comparisons across the entire brain ($p<0.005$). Note that although the effects resulting from the suboptimal 5 TR control analysis are weaker, they are still anatomically similar to the effects resulting from the 10 TR control analysis.

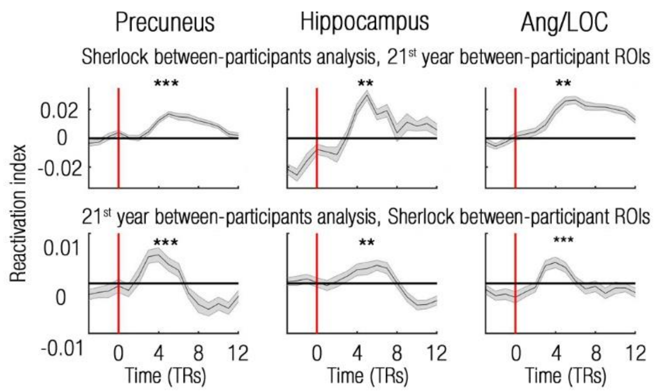


**Supplementary Figure 5. Reactivation is specific to event-boundaries – replication across datasets using a scene-based between-participant analysis.**
Reactivation indices are presented for the time period of 3 TRs prior to event boundaries until 12 TRs after event boundaries, uncorrected for the HRF delay. Event boundaries are depicted as red vertical lines. Time-courses are presented for the bilateral precuneus/retrosplential cortex (PCUN), hippocampus and Angular gyrus/Lateral Occipital Cortex (Ang/LOC) ROIs (supplementary Figure 2). ROIs were defined based on one dataset (Sherlock/21st year) and tested on the other dataset using the scene-based between participant analysis. ** p<0.01; *** p<0.001.

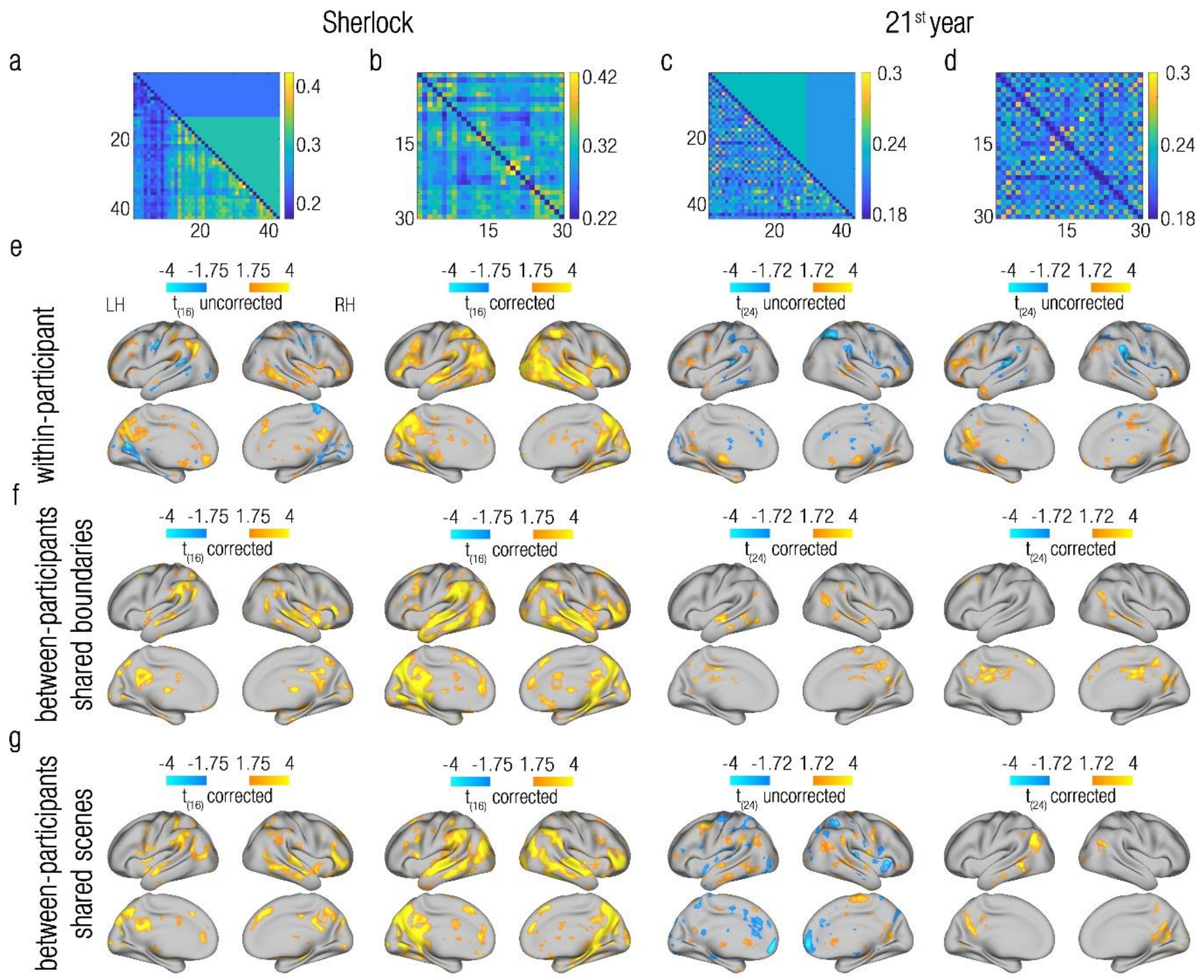


**Supplementary Figure 6. Removal of time-related biases from the Bag Of Words analyses.**
**(a)** Bag of Words scene-similarity matrix of the Sherlock movie. The upper triangular part of the matrix holds the mean of the corresponding symmetrical entries in the lower triangular part of the matrix. Note the difference in means between the first 15 scenes and the rest of the scenes. **(b)** The same matrix presented in (a), after removal of the first 15 scenes. **(c)** Bag of Words scene-similarity matrix of the 21$^{st}$ year story. The upper triangular part of the matrix holds the mean of the corresponding congruent symmetrical entries in the lower triangular part of the matrix. Note the difference in means between the last 15 scenes and the rest of the scenes. **(d)** The same matrix presented in (c), after removal of the last 15 scenes. **(e)** Within-participant Bag of Words analysis, using each of the matrices presented in (a-d) and their related fMRI dataset. **(f)** Boundary-based between-participant Bag of Words analysis, using each of the matrices presented in (a-d) and their related fMRI dataset. **(g)** Scene-based between-participant Bag of Words analysis, using each of the matrices presented in (a-d) and their related fMRI dataset. Note that although the effects related to the time-biased matrices are weaker (some not surviving correction for multiple comparisons, as indicated by the corresponding colour-bars), they are still anatomically similar to the effects related to the unbiased matrices.

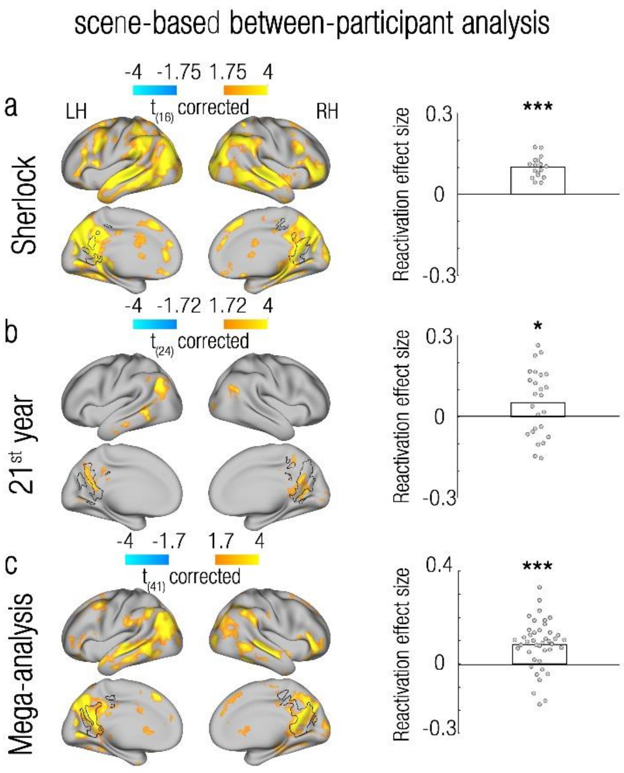


**Supplementary Figure 7. Relevant past events are preferentially reactivated at event boundaries – scene-based between-participant analysis.**

The level to which reactivation is modulated by semantic context (as defined by a Bag Of Words model, Figure 1d) was measured across the entire brain. Rows show within-dataset results of the Sherlock dataset **(a)**, the 21$^{st}$ year dataset **(b)**, and a mega-analysis pooling together the two datasets **(c)**. All maps were cluster-corrected for multiple comparisons across the entire brain ($p<0.005$). Maps are projected onto a flat cortical surface. Similarly to the results of the within-participants analysis and the boundary-based between participant analysis (Figure 5), the precuneus/retrosplenial cortex showed a consistent positive modulation of reactivation by semantic context across datasets (scenes with semantic context similar to that of each event boundary were reactivated more than scenes with different semantic contexts). Independent precuneus/retrosplenial cortex ROIs derived from each dataset (Sherlock/21$^{st}$ year) in the scene-based between-participant reactivation analysis (Supplementary Figure 2) were used to test the reactivation-modulation effects in the same dataset and using the same analysis method (ROIs depicted in black contours). Group means are represented as bars, and single-participant scores are represented as circles. This ROI analysis confirmed that the same region that showed significant reactivation effects also showed positive modulation of reactivation by sematic context. LH, left hemisphere; RH, right hemisphere; * $p<0.05$; *** $p<0.001$.

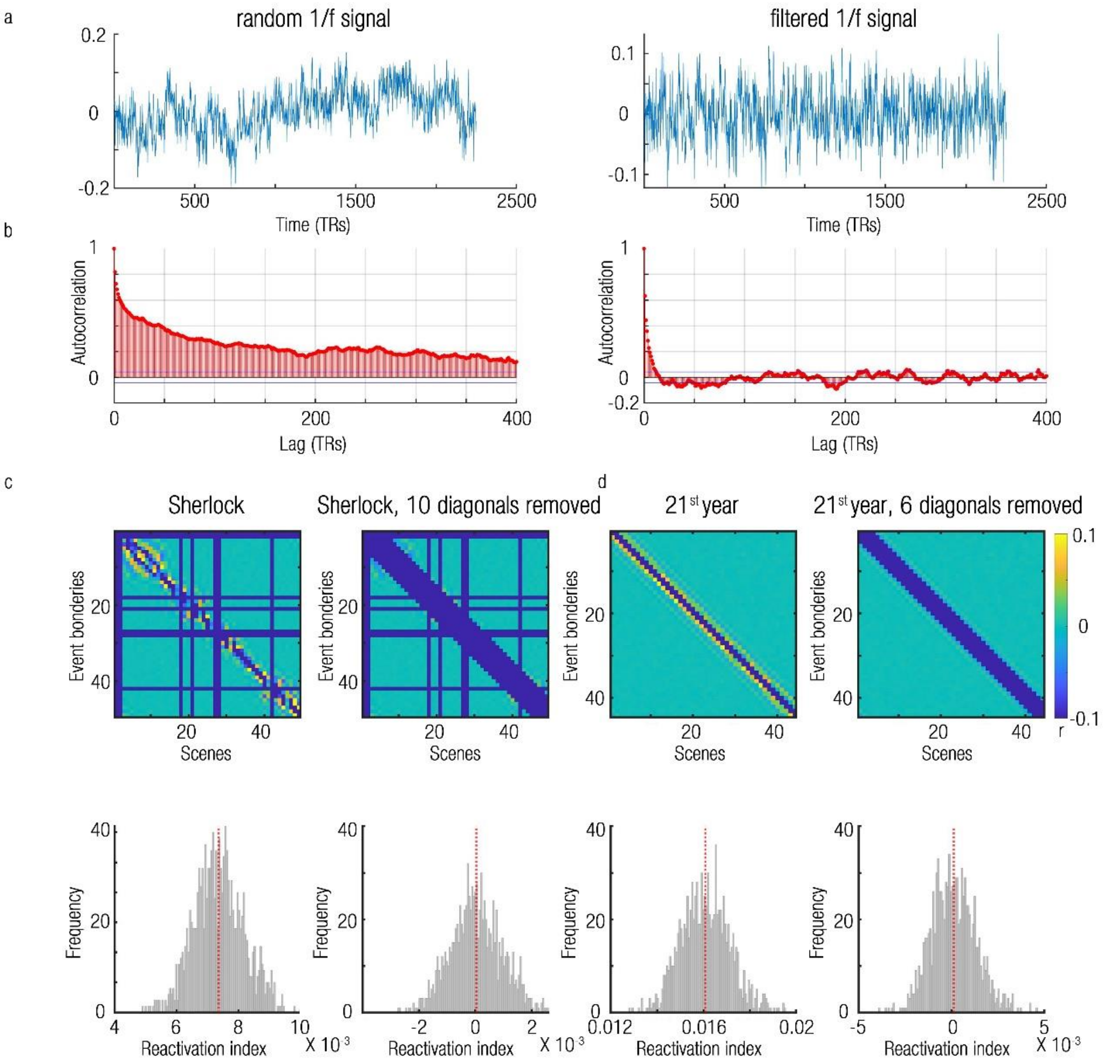


**Supplementary Figure 8. Removal of temporal filtering artefacts from the within-participant reactivation analysis.**

**(a)** Left: random signal with a 1/f power-spectrum. Right: the same signal after the application of high-pass filtering. Note the removal of slow modulations in the signal. **(b)** Left: the autocorrelation structure of the unfiltered signal presented in (a). Right: the autocorrelation structure of the filtered signal presented in (a). Note the alternating positive and negative autocorrelations that diminish across time. **(c)** Upper left: averaged event boundary X scene similarity matrix, based on filtered 1/f signals, simulated according to characteristic of the Sherlock dataset. Blue rows/columns correspond to excluded scenes and event boundaries (see Online Methods). Note the artefactual correlations around the main diagonal. These correlations result in a biased distribution of the reactivation indices (bottom left, mean depicted with a vertical dashed red line). Upper right: removal of 10 diagonals from the simulated matrices masks out the artefactual correlations. This procedure results in an unbiased distribution of the reactivation indices (bottom right, mean is centred on zero, as depicted with a vertical dashed red line). **(d)** The same simulation analysis as in (c), but this time based of the characteristics of the 21st year dataset. Artefactual correlations and the resulting biased distribution of the reactivation indices (left) can be corrected by removing 6 diagonals of the similarity matrices, thus centring the distribution of the reactivation indices on zero (right). Note that the temporal extent of the artefact (and hence the required amount of removed diagonals) depends on the specific characteristics of each dataset, such as the lengths of scenes, their temporal arrangement and the width of the high-pass filter employed.

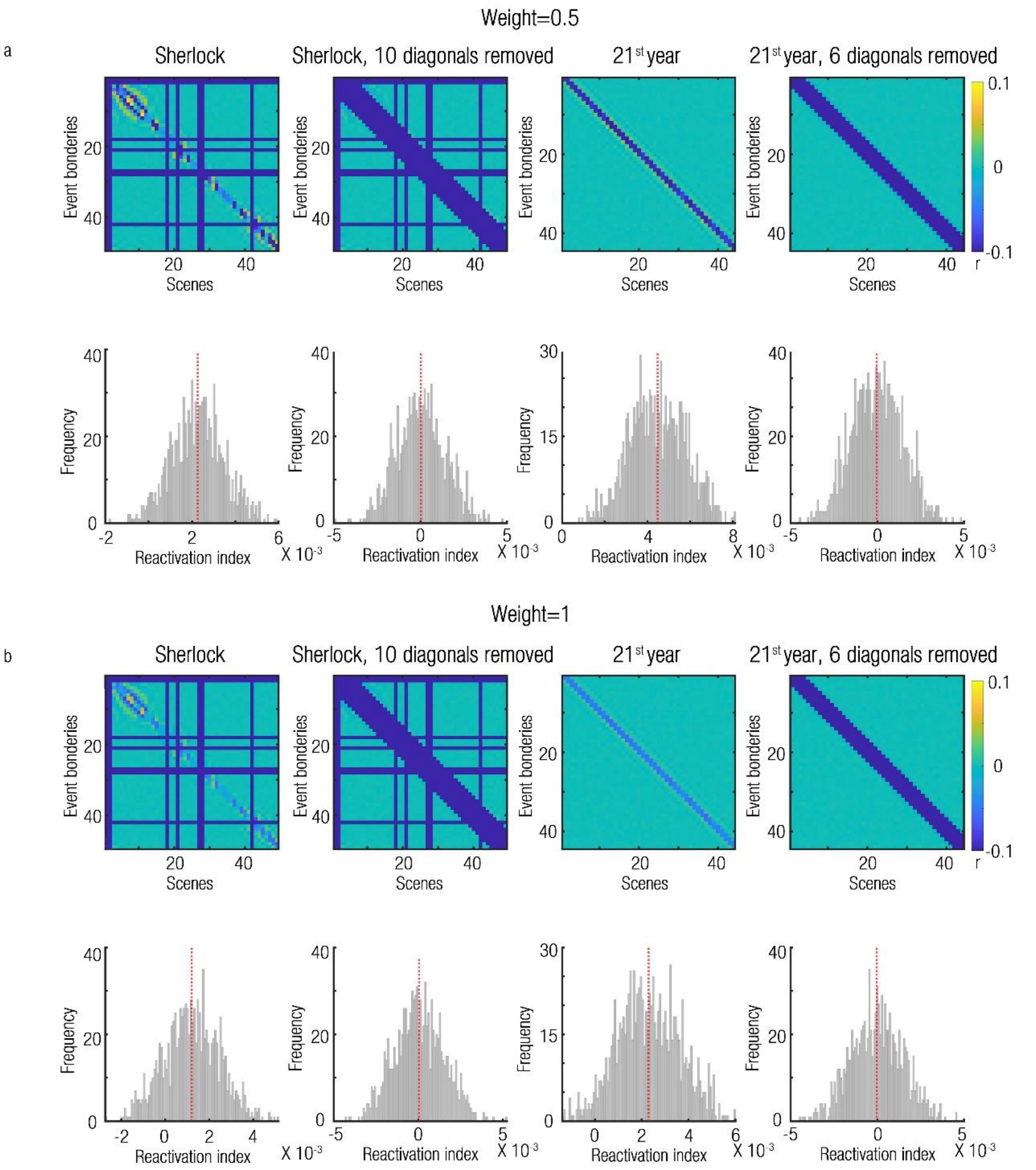


**Supplementary Figure 9. Simulation of constant scene representations in the within-participant reactivation analysis.**

Same as sup. Figure 8c-d, but this time random scene representations have been added in with a weight of 0.5 (a) or 1 (b). Note that the temporal filtering artefact was reduced with increased weights, but still remained, and required the removal of diagonals from the analysis.

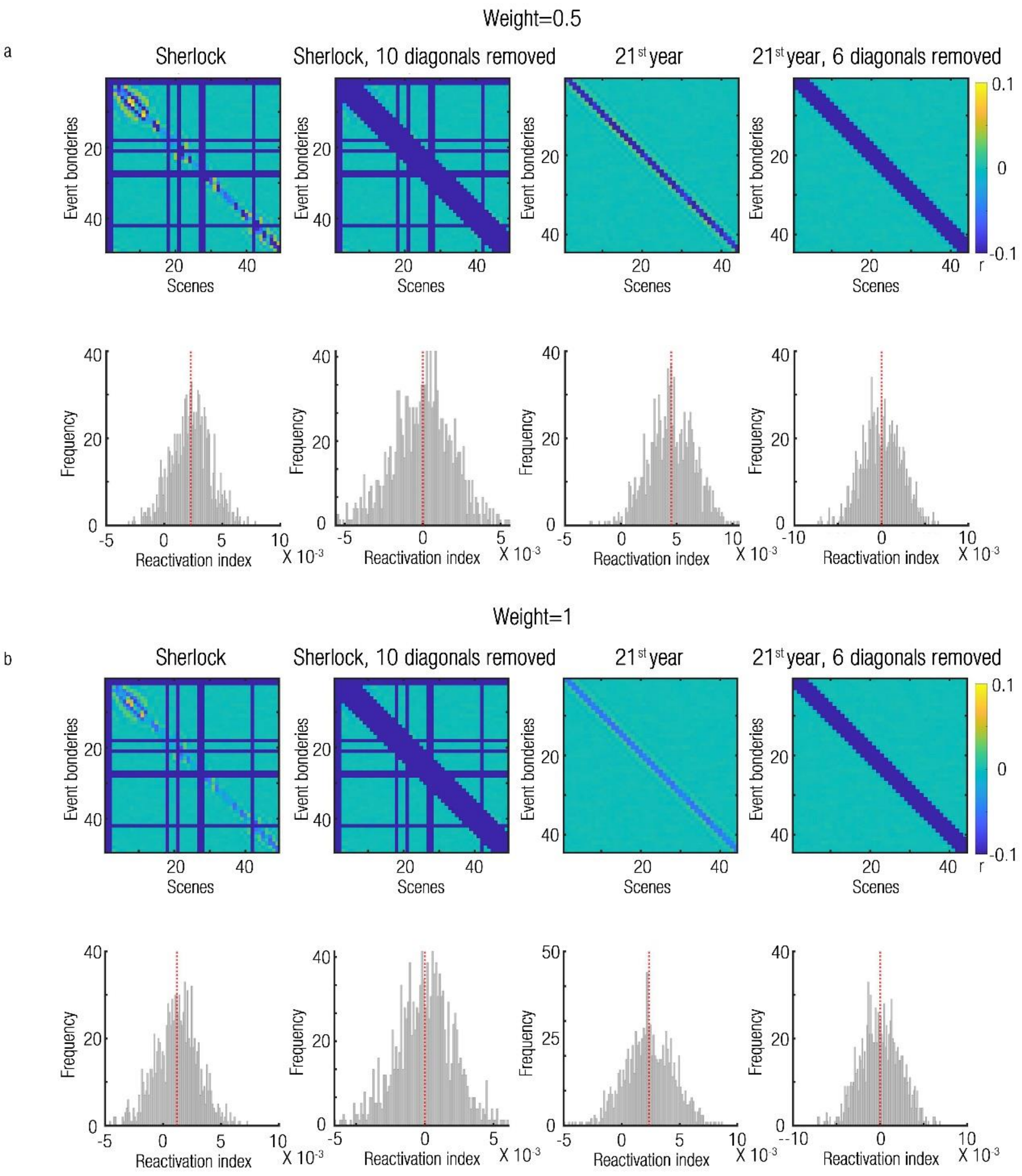


**Supplementary Figure 10. Simulation of autocorrelated scene representations in the within-participant reactivation analysis**

Same as sup. Figure 8c-d, but this time autocorrelated random scene representations have been added in with a weight of 0.5 (a) or 1 (b). Note that the temporal filtering artefact was reduced with increased weights, but still remained, and required the removal of diagonals from the analysis.

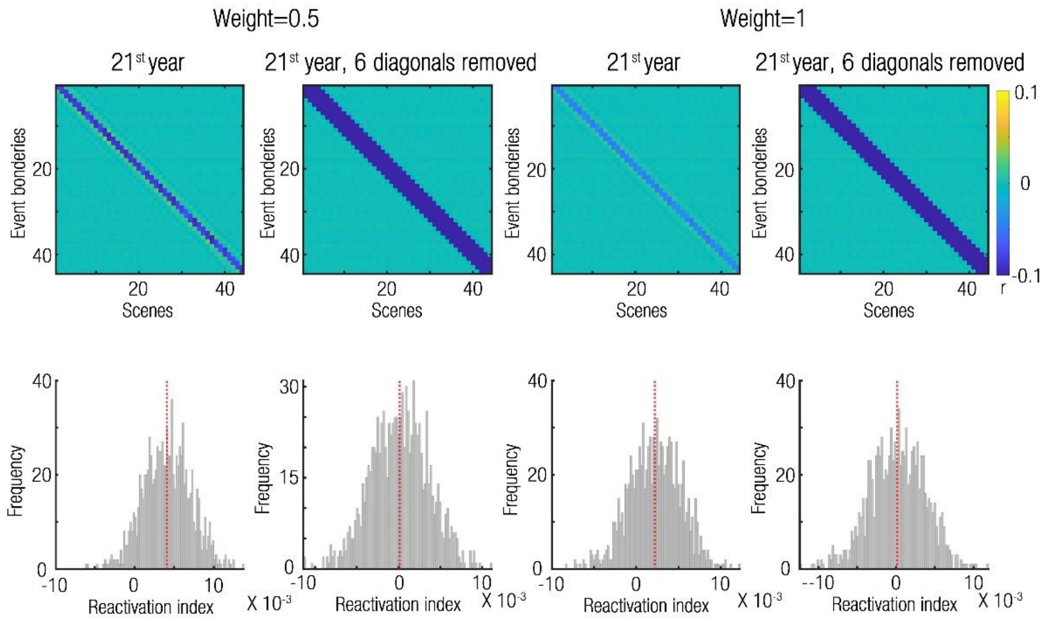


**Supplementary Figure 11. Simulation of congruency in scene representations in the within-participant reactivation analysis of the 21st year dataset.**
Same as sup. Figure 8d, but this time two series of correlated random scene representations have been added in an interleaved manner, with a weight of 0.5 (left) or 1 (right), such that a congruency pattern was formed. Note that the temporal filtering artefact was reduced with increased weights, but still remained, and required the removal of diagonals from the analysis.

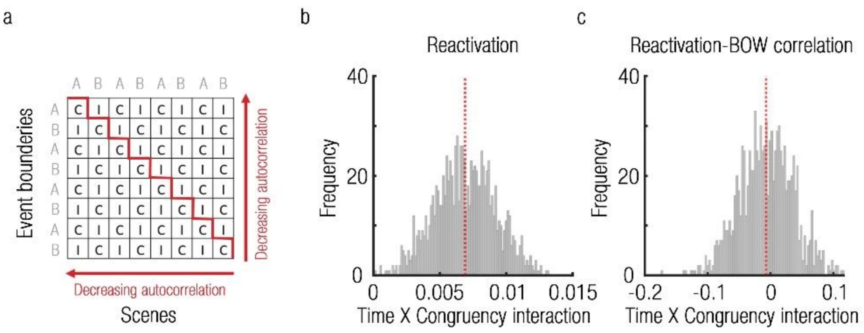


**Supplementary Figure 12. Congruency-related artefacts induced by the alternating narrative structure of the 21st year dataset.**

**(a)** The first 30 scenes of the 21st year dataset consist of alternating narratives (A/B, marked in grey). The similarity matrices calculated for this dataset (schematically depicted here) would therefore contain alternating entries of congruent (same narrative, denoted with C) or incongruent (different narrative, denoted with I) correlations between event boundaries and scenes. These entries are arranged by diagonals, which create an asymmetric congruency structure with regards to event boundaries. For example, the main diagonal contains correlations between event boundaries and their immediately preceding scenes, which are congruent. The superdiagonal contains correlations between event boundaries and their immediately following scenes, which are incongruent. However, autocorrelations have a symmetrical effect in time (illustrated by red arrows). Event boundaries are more correlated to closer points in time, and are hence more correlated to congruent scenes in the past and to incongruent scenes in the future, which will bias any congruency-related analysis. This bias cannot be fixed by removing diagonals from the analysis, since an equal number of diagonals must be removed from the past/future parts of the matrix, which will maintain an asymmetrical congruency structure in time. **(b)** Simulation analysis, using filtered 1/f signals, simulated according to characteristic of the 21st year dataset (see Online Methods). When defining a reactivation index as the Time (past/future) X Congruency (same/different narrative) interaction, the distribution of reactivation indices remains biased despite the removal of 6 diagonals (compare with sup. Figure 8d). **(c)** Simulation analysis, using filtered 1/f signals, simulated according to characteristic of the 21st year dataset. These signals were correlated with the 21st year Bag of Words (BOW) scene-similarity matrix, such that a Time X Congruency interaction of correlations was computed (see Online Methods). This resulted in a negative bias in the distribution of interaction values. Dashed red lines mark distribution means.

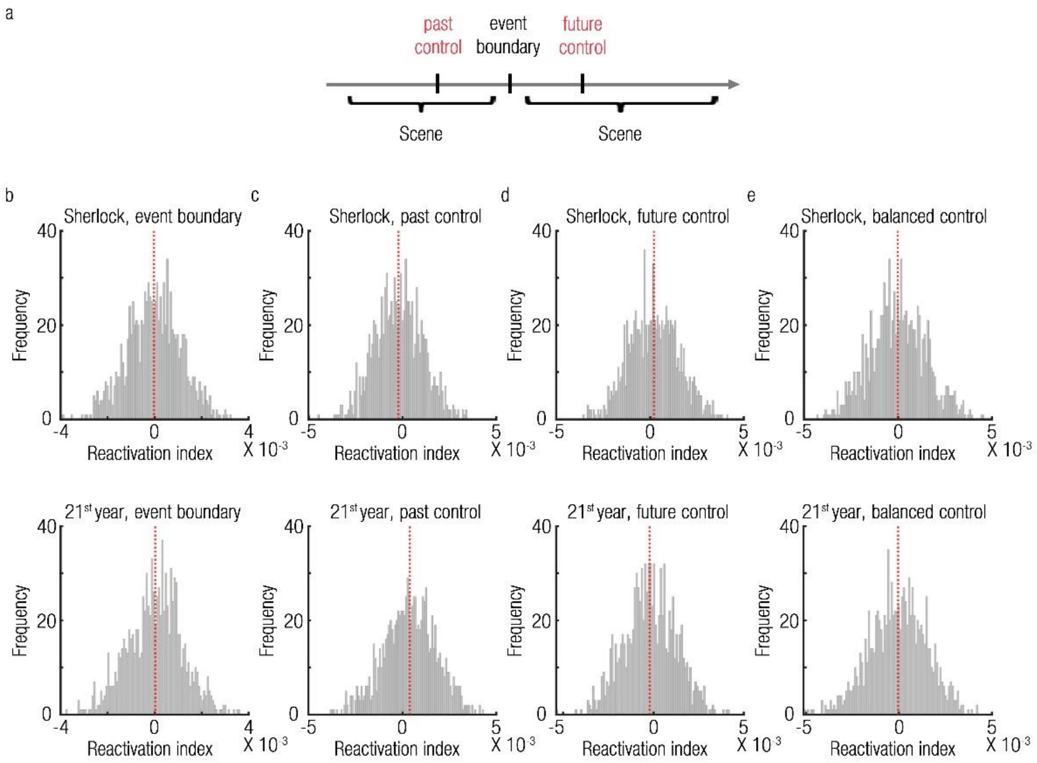


**Supplementary Figure 13. Simulation of the controlled reactivation analysis.**
**(a)** Illustration of a past control timepoint and a future control timepoint, both equally distant from each event boundary. **(b)** The distributions of reactivation indices at event boundaries, derived from random signals with the characteristics of the Sherlock (top) or the 21st year (bottom) datasets, were centered on zero after removal of the relevant number of diagonals (as in sup. Figure 8c/d). **(c)** The distributions of reactivation indices at the past control points showed biases in both simulated datasets. **(d)** The distributions of reactivation indices at the future control points also showed biases in both simulated datasets. **(e)** A balanced control was defined as the average reactivation index across the past and future timepoints. The contrast between the reactivation indices at event boundaries and those of the balanced controls resulted in unbiased distributions that were centered on zero. Dashed red lines mark distribution means.
